# Fathers’ preconception smoking and offspring DNA methylation: A two generation study

**DOI:** 10.1101/2023.01.13.523912

**Authors:** Negusse T Kitaba, Gerd Toril Mørkve Knudsen, Ane Johannessen, Faisal I. Rezwan, Andrei Malinovschi, Anna Oudin, Bryndis Benediktsdottir, David Martino, Francisco Javier Callejas González, Leopoldo Palacios Gómez, Mathias Holm, Nils Oskar Jõgi, Shyamali C Dharmage, Svein Magne Skulstad, Sarah H Watkins, Matthew Suderman, Francisco Gómez-Real, Vivi Schlünssen, Cecilie Svanes, John W. Holloway

**Author notes:** **Correspondence:** Name: Prof John W. Holloway, Address: Human Development and Health, Faculty of Medicine, University of Southampton, Southampton SO16 6YD, UK. Joint first authors (Negusse T Kitaba, Gerd Toril Mørkve Knudsen). Joint senior authors (Cecilie Svanes, John W. Holloway). **Funding:** Co-ordination of the RHINESSA study has received funding from the Research Council of Norway (Grants No. 274767, 214123, 228174, 230827 and 273838), ERC StG project BRuSH #804199, the European Union’s Horizon 2020 research and innovation programme under grant agreement No. 633212 (the ALEC Study), the Bergen Medical Research Foundation, and the Western Norwegian Regional Health Authorities (Grants No. 912011, 911892 and 911631). Study centres have further received local funding from the following: Bergen: the above grants for study establishment and co-ordination, and, in addition, World University Network (REF and Sustainability grants), Norwegian Labour Inspection, and the Norwegian Asthma and Allergy Association. Albacete and Huelva: Sociedad Española de Patología Respiratoria (SEPAR) Fondo de Investigación Sanitaria (FIS PS09). Gøteborg, Umeå and Uppsala: the Swedish Heart and Lung Foundation, the Swedish Asthma and Allergy Association. Reykjavik: Iceland University. Melbourne: National Health and Medical Research Council (NHMRC) of Australia (research grants 299901 and 1021275). Tartu: the Estonian Research Council (Grant No. PUT562). Århus: The Danish Wood Foundation (Grant No. 444508795), the Danish Working Environment Authority (Grant No. 20150067134), Aarhus University (PhD scholarship). ALSPAC funding: EPIC arrays age 15-17, John Templeton Foundation (60828) and EPIC arrays age 24, MRC (MC_UU_12013/2) & CLOSER (MRC and ESRC). **Author Contributions** Conceptualization: Cecilie Svanes, Gerd Toril Mørkve Knudsen, Faisal I Rezwan, Ane Johannessen, Negusse Kitaba, John W Holloway Data curation: Gerd Toril Mørkve Knudsen, Bente Skottvoll, Ane Johannessen, Negusse Kitaba, John W Holloway Formal analysis: Negusse Kitaba, Cecilie Svanes, John W Holloway Methodology: Negusse Kitaba, Faisal I Rezwan, Cecilie Svanes, John W Holloway Replication: Sarah Watkins, Matthew Suderman Project administration: Cecilie Svanes Writing – original draft: Negusse Kitaba, Gerd Toril Mørkve Knudsen, Cecilie Svanes, John W Holloway Writing – review, editing and final approval: All authors.

## Abstract

**Rationale:** Experimental studies suggest that exposures may impact respiratory health across generations via epigenetic changes transmitted specifically through male germ cells. Studies in humans are however limited. We aim to identify epigenetic marks in offspring associated with father’s preconception smoking.

**Methods:** We conducted epigenome-wide association studies (EWAS) in the RHINESSA cohort on father’s any preconception smoking (N=875 offspring) and father’s pubertal onset smoking <15 years (N=304), using Infinium MethylationEPIC Beadchip arrays, adjusting for offspring age, maternal smoking and personal smoking. EWAS of maternal and offspring personal smoking were performed for replication.

**Results:** Father’s smoking commencing preconception was associated with methylation of blood DNA in offspring at two Cytosine-phosphate-Guanine sites (CpGs) (False Discovery Rate (FDR) <0.05) in *PRR5* and *CENPP*. Father’s pubertal onset smoking was associated with 19 CpGs (FDR <0.05) mapped to 14 genes (*TLR9, DNTT, FAM53B, NCAPG2, PSTPIP2, MBIP, C2orf39, NTRK2, DNAJC14, CDO1, PRAP1, TPCN1, IRS1* and *CSF1R*). These differentially methylated sites were hypermethylated and associated with promoter regions capable of gene silencing. Some of these sites were associated with offspring outcomes in this cohort including ever-asthma (NTRK2), ever-wheezing (DNAJC14, TPCN1), weight (FAM53B, NTRK2) and BMI (FAM53B, NTRK2) (P< 0.05). Pathway analysis showed enrichment for gene ontology pathways including regulation of gene expression, inflammation and innate immune responses.

**Conclusion:** Father’s preconception smoking, particularly in puberty, is associated with offspring DNA methylation, providing evidence that epigenetic mechanisms may underly epidemiological observations that pubertal paternal smoking increases risk of offspring asthma, low lung function and obesity.

## Introduction

There is growing consensus that perturbations of the epigenome through parental exposures even before their offspring are conceived may explain some of the variation in the heritability of health and disease not captured by Genome-Wide Association Studies (GWAS). The period of puberty in future parents, in particular fathers, may represent a critical window of physiological change and epigenetic reprogramming events, which may increase the individual’s susceptibility for environmental exposures to be embodied in the developing gametes^1,2^. Animal and human studies have shown that prenatal as well as personal exposure to smoking are associated with epigenetic modifications that impact on sperm count and quality^3^. There is now growing interest in how epigenetic modifications, such as DNA methylation (DNAm), related to the parental *preconception* period may influence the health of the *next generation*^4^.

Although smoking rates are generally declining, smoking commencing before the age of 15 is increasing^5,6^. Epidemiological studies have demonstrated that father’s smoking in adolescent years may be a causal factor for poorer respiratory health in offspring. Both fathers’ smoking initiation before age 15 and smoking duration before conception have been associated with more asthma and lower lung function in offspring^7–9^. Father’s preconception smoking onset has also been associated with higher body fat mass in sons^10–13^.

Epigenome-Wide Association Studies (EWAS) have identified extensive methylation biomarkers associated with personal smoking, all-cause mortality in current and former smokers, as well as mother’s smoking during pregnancy^6,14–17^. While previous studies have identified DNA methylation signals in offspring blood^18^ and cord blood^19^ related to father’s smoking, they have not specifically investigated the timing of exposure, partly because detailed smoking information from fathers is rarely available^20^. Methylation markers associated with paternal preconception smoking, could have an important role in elucidating long-term effects on the offspring epigenome, with the potential for developing efficient intervention programs and improved public health.

This study aimed to investigate whether DNA methylation of DNA measured in offspring blood is associated with fathers’ smoking commencing before conception, and in particular, with fathers’ smoking starting in (pre)pubertal years (before age 15). We hypothesized that epigenetic changes involving DNA methylation may explain the molecular mechanisms underlying the association between fathers’ smoking preconception and offspring health observed in epidemiological studies. Additionally, we hypothesized that fathers’ smoking in the critical window of early puberty may have a more significant impact on the offspring epigenome. In a two-generation cohort, we sought to identify the DNA methylation changes in offspring blood associated with (1) father’s smoking onset preconception compared with never or later onset smoking, and (2) father’s smoking onset before age 15 compared with never smoking.

## Methods

### Study design and data

We used data and samples from offspring that participated in the RHINESSA study (www.rhinessa.net). Parent data, including detailed information on smoking habits, were retrieved from the population-based European Community Respiratory Health Survey (ECRHS, www.ecrhs.org) and/or the Respiratory Health in Northern Europe study (RHINE, www.rhine.nu) studies. This analysis comprised 875 offspring-parent pairs with complete information, from six study centres with available peripheral blood for offspring (Aarhus, Denmark; Albacete/Huelva, Spain; Bergen, Norway; Melbourne, Australia; Tartu, Estonia). All participants were of Caucasian ancestry. Medical research committees in each study centre approved the studies, and each participant gave written consent.

Father’s smoking and age of starting/quitting was reported in interviews/questionnaires, and related to offspring’s birth year, to define the categories: never smoked (N=547), any preconception smoking (N=328), preconception smoking with onset <15 years (pubertal smoking) (N=64) (cut point based on mean age of voice break 14.5 years, first nocturnal seminal emission 14.8 years). Personal smoking was classified as current, ex- or never smoking. Maternal smoking was defined by offspring’s report on mothers’ smoking during their childhood/pregnancy.

DNAm in offspring was measured using Illumina Infinium MethylationEPIC Beadchip arrays (Illumina, Inc. CA, USA) and data processed using an established pipeline as detailed in the online supplement. Following processing 726,661 CpGs were retained for analysis.

### Statistical analysis

Two EWAS on preconception paternal smoking as exposure (any preconception smoking, prepuberty smoking) using robust regression were run with offspring blood DNA methylation as outcome adjusting for offspring’s sex, age, personal and mother’s smoking, study center and cell-type proportions at significance level of false discovery rate (FDR) corrected p-value <0.05. Inflation from systematic biases was adjusted using BACON. Differentially methylated regions were detected using DMRCate and dmrff. In additional analyses, associations between fathers’ any preconception smoking and offspring’s DNA methylation were also stratified by offspring sex. Biological interpretation of significant dmCpGs is detailed in the supplementary methods.

We compared our EWAS results with findings from meta-analyses of EPIC array DNA methylation associated with personal smoking from four population-based cohorts^21^, personal smoking-methylation effects from 16 cohorts using 450K arrays^16^; and the Pregnancy and Childhood Epigenetics Consortium (PACE) meta-analysis of mother’s smoking in pregnancy on offspring cord blood methylation^22^ to assess the shared count of dmCpG sites at (FDR<0.05) for the overlap between each EWAS.

#### Replication analysis

Replication was carried out in the ALSPAC (Avon Longitudinal Study of Parents and Children) cohort adjusted for predicted cell count proportions, batch effects (plate), maternal smoking during pregnancy, self-reported own smoking, age and sex using DNA methylation data from whole blood measured at age 15-17. A description of the ALSPAC cohort is provided in the supplementary methods. T-tests were used to compare the association of regression coefficient of RHINESSA’s dmCpG sites at FDR <0.05 and the top 100 CpG sites with ALSPAC. Signed tests were used to test the direction of association.

### Sensitivity analyses

To assess the effect of social class, father’s education was used as a proxy for social class. In order to see the effect of CpGs changing with age, the correlation of methylation at dmCpGs known to be associated with offspring age, known aging markers from RHINESSA EWAS, dmCpG sites for father smoking before age 15, and offspring age was assessed. To further investigate whether the identified dmCpGs were associated with respiratory outcomes and weight in the offspring, we conducted regression analysis between offspring’s DNA methylation signals and offspring’s own reports of ever-asthma, ever-wheeze, weight and BMI, while accounting for offspring sex

## Results

The analysis included 875 RHINESSA participants (Table 1A), 457 males and 418 females, aged 7 to 50 years. Of these 328 had a father who had ever smoked of which 64 had started before age 15 years; 263 had a mother who had ever smoked, and 240 had smoked themselves. Characteristics are also given for the sub-sample of 304 offspring whose father either had started smoking before age 15 years, or never smoked (Table 1B).

**Table 1 A and B:** General characteristics of study participants from the RHINESSA study with complete data on offspring DNA methylation and father’s age of onset of tobacco smoking. A: for the full cohort of 875 offspring, and B: for the 304 offspring whose father started to smoke before age 15 years or never smoked.

| Characteristic | A: FULL Cohort |  |  | B: Start Smoking <15 years |  |  |
| --- | --- | --- | --- | --- | --- | --- |
|  | No, N = 547<sup>1</sup> | Yes, N = 328<sup>1</sup> | p-value<sup>2</sup> | No, N = 240<sup>1</sup> | Yes, N = 64<sup>1</sup> | p-value<sup>3</sup> |
| **Sex** |  |  | 0.8 |  |  | 0.5 |
| F | 263 (48%) | 155 (47%) |  | 112 (47%) | 33 (52%) |  |
| M | 284 (52%) | 173 (53%) |  | 128 (53%) | 31 (48%) |  |
| **Study centre** |  |  | <0.001 |  |  | 0.005 |
| Albacete | 24 (4.4%) | 32 (9.8%) |  | 7 (2.9%) | 9 (14%) |  |
| Arhus | 34 (6.2%) | 17 (5.2%) |  | 14 (5.8%) | 1 (1.6%) |  |
| Bergen | 320 (59%) | 194 (59%) |  | 174 (72%) | 47 (73%) |  |
| Huelva | 17 (3.1%) | 14 (4.3%) |  | 5 (2.1%) | 2 (3.1%) |  |
| Melbourne | 78 (14%) | 14 (4.3%) |  | 21 (8.8%) | 1 (1.6%) |  |
| Tartu | 74 (14%) | 57 (17%) |  | 19 (7.9%) | 4 (6.2%) |  |
| **Age** | 26 (8) | 30 (8) | <0.001 | 26 (8) | 27 (8) | 0.2 |
| **Mother smoking** | 84 (15%) | 179 (55%) | <0.001 | 30 (12%) | 34 (53%) | <0.001 |
| **Offspring Smoking** | 121 (22%) | 119 (36%) | <0.001 | 44 (18%) | 25 (39%) | <0.001 |
| **B-cells** | 0.022 (0.019) | 0.020 (0.016) | 0.4 | 0.022 (0.016) | 0.020 (0.015) | 0.4 |
| **CD4-cells** | 0.030 (0.031) | 0.032 (0.032) | 0.3 | 0.026 (0.026) | 0.026 (0.024) | >0.9 |
| **CD8-cells** | 0.13 (0.05) | 0.12 (0.05) | 0.5 | 0.13 (0.04) | 0.12 (0.04) | 0.7 |
| **NK-cells** | 0.07 (0.04) | 0.07 (0.04) | 0.8 | 0.07 (0.04) | 0.07 (0.04) | 0.4 |
| **Mononuclear cells** | 0.073 (0.022) | 0.071 (0.020) | 0.2 | 0.074 (0.020) | 0.071 (0.020) | 0.4 |
| **Neutrophil** | 0.67 (0.09) | 0.68 (0.08) | 0.3 | 0.68 (0.08) | 0.68 (0.07) | 0.8 |

<sup>1</sup> n (%); Mean (SD) <sup>2</sup> Pearson's Chi-squared test; Wilcoxon rank sum test <sup>3</sup> Pearson's Chi-squared test; Fisher's exact test; Wilcoxon rank sum test
\*Including father never smoked and father started smoking after birth of the offspring

### Epigenome wide association analysis of preconception father’s smoking

Epigenome-wide association between father’s any preconception smoking and offspring DNA methylation identified two dmCpGs (inflation λ=1.187); cg00870527 mapped to *PRR5* and cg08541349 mapped to *CENPP* (Table 2A, and supplementary table E1). The genome-wide distribution of associated dmCpGs is shown in Figure 1A. The comparison of methylation distribution between never- and ever-smoke exposed is shown in Figure 1C.

**Table 2A and B.**
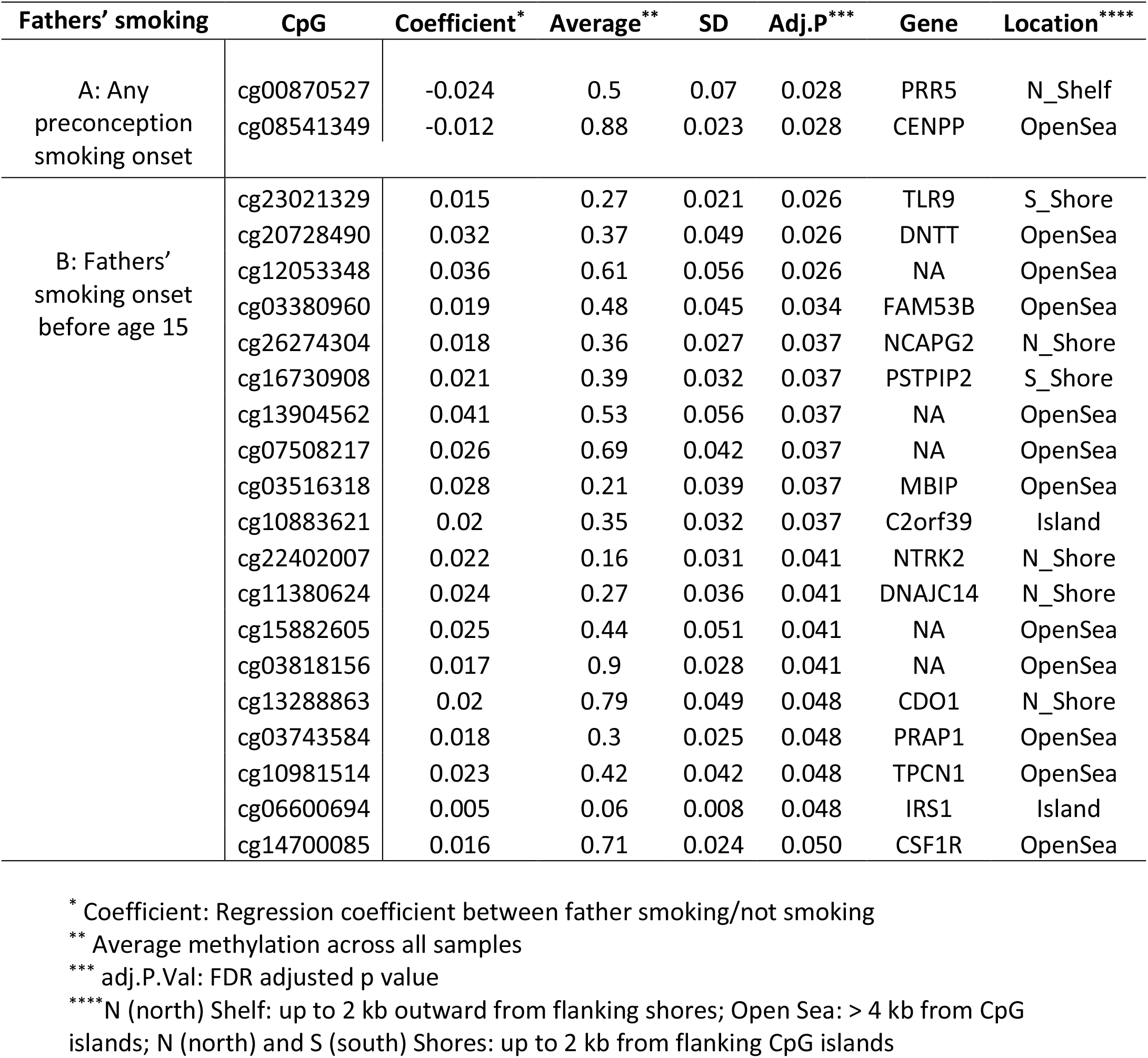
CpG sites associated with father’s smoking at genome wide significance (FDR<0.05) **A:** for father’s any preconception smoking, in the full cohort (N=875), and **B**: for father’s smoking starting before age 15 years, in the subpopulation (N=304).

**Figure 1A and B.**
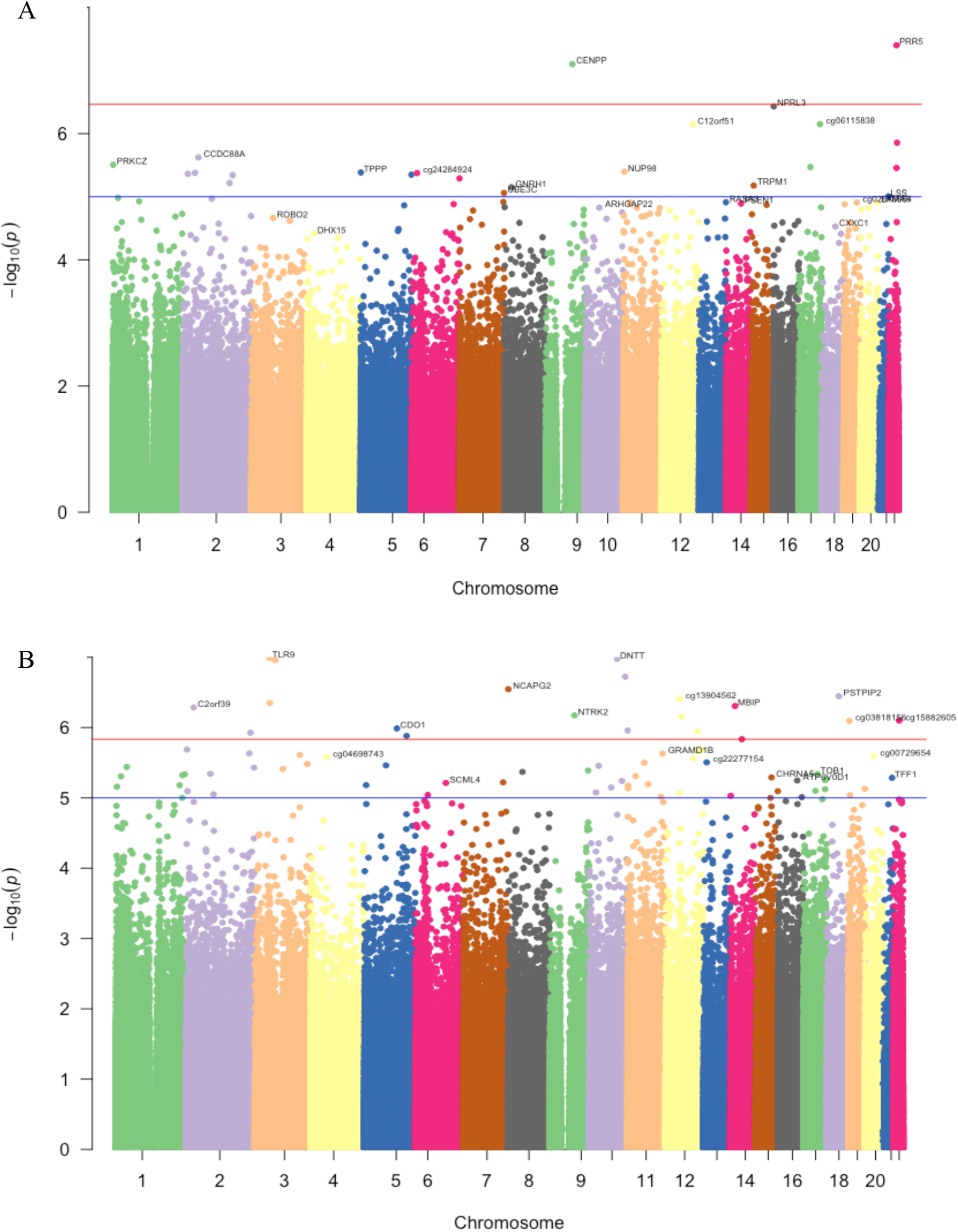
Manhattan plot for Genome-wide distribution of dmCpGs **A:** for father’s any preconception smoking, and **B:** father’s pubertal smoking starting before age 15. The red line shows genome-wide significance, the blue is the suggestive line. The y-axis represents -log10 of the p-value for each dmCpG (indicated by dots) showing the strength of association. The x-axis shows the position across autosomal chromosomes. The top dmCpGs on each chromosome were annotated to the closest gene.

**Figure 1C and D.**
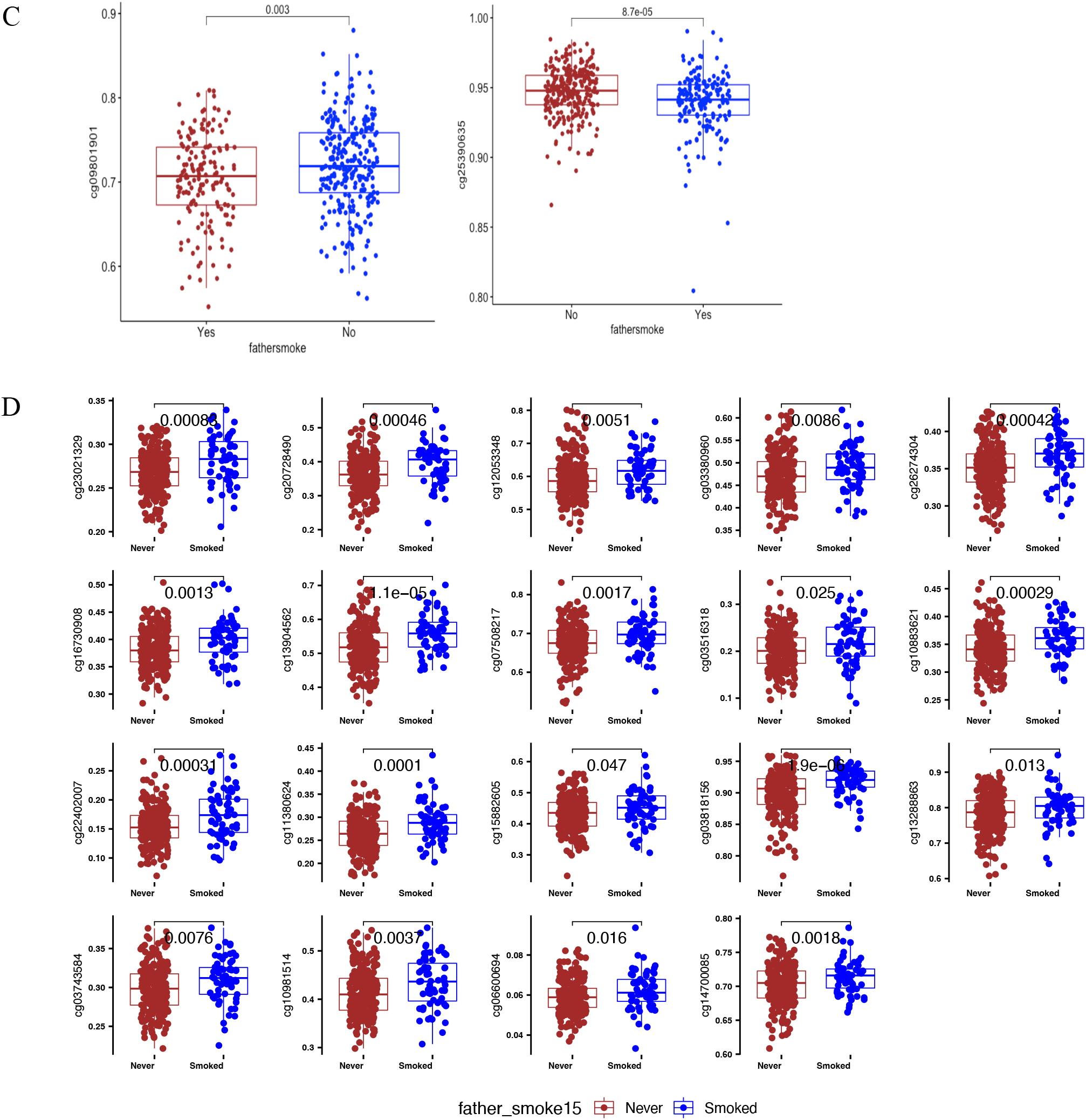
Comparison of methylation differences for **C**: for father’s any preconception smoking, and **D**: for father’s pubertal smoking starting before age 15.

In sex-stratified analysis, in males (N=457) we identified four dmCpGs mapped to KCNJ1, GRAMD4, TRIM2 and MYADML2. In females (N=418) there was one dmCpG mapped to LEPROT1 (FDR <=0.05) (Supplementary Table E2). All sex-specific dmCpGs were hypomethylated.

To specifically determine the signature related to father’s early onset smoking, we compared methylation differences between offspring of fathers who started to smoke <15 years (n=64) with offspring of never smoking fathers (n=240). We identified 55 dmCpGs at FDR <0.05 (λ=1.44) showing genome-wide significance. After adjusting for inflation using BACON, 19 dmCpGs showed significant association at FDR <0.05 with λ=1.29 (Table 2B, Figure 1B, and supplementary Table E3). These dmCpGs were mapped to 14 known genes and 5 intergenic regions. The genes include TLR9, DNTT, FAM35B, NCAPG2, MBIP, C2orf39, NTRK2, DNAJC14, CDO1, PRAP1, TPCN1, IRS1, PSTPIP2, and CF1R. All hits were hypermethylated in the exposed group. The comparison of methylation distribution between the never and smoke exposed is shown in Figure 1D.

The dmCpGs associated with father’s preconception smoking were mainly located in open-sea genomic features and enriched for promoter regions (Table 2A). The dmCpGs associated with father’s pubertal smoking were in open-sea genomic features and CpG island shores (flanking shore regions, <2 kb up-and downstream of CpG islands) and enriched for CpG islands and gene bodies (Table 2B).

### Father’s preconception smoking signatures as compared with signatures of personal and mother’s smoking

To compare the effects of father’s preconception and pubertal smoking on the offspring epigenome with that of other smoking exposures, the epigenome-wide effects of offspring’s own smoking as well as their mother’s smoking during pregnancy and childhood were assessed. We identified 33 dmCpGs related to personal smoking, and 14 dmCpGs associated with mother’s smoking (FDR<0.05) (Supplementary Tables E4 and E5, respectively).

To illustrate the distinct and shared genome-wide effects of personal, mother’s, and father’s smoking on the offspring methylome, we generated a locus-by-locus genome comparison, (Figure 2A). While there was similarity between the effects of personal smoking and mother’s smoking on chromosome 5, we observed distinct signatures for father’s preconception smoking on chromosome 22, and for mother’s smoking exposure on chromosomes 7 and 15.

**Figure 2A.**
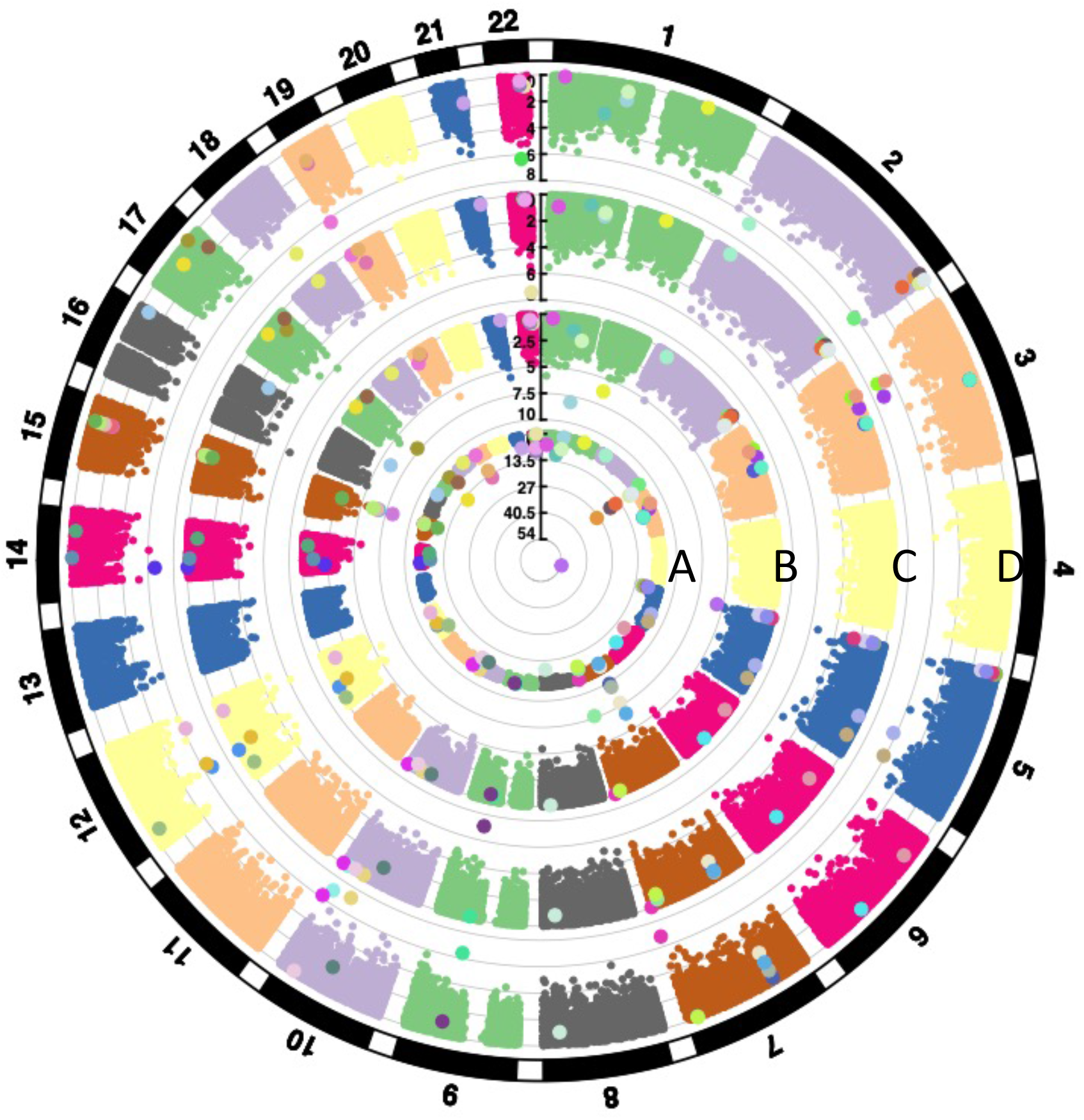
Circos plots showing genome-wide distribution across autosomal chromosomes of dmCpGs associated with **A:** personal smoking (in offspring), **B:** mother’s smoking, **C:** father’s any preconception smoking, and **D:** father’s pubertal smoking starting before age 15. Each dot represents a CpG site; the radial line shows the -log10 p-value for each EWAS. Zoomed dots show significant sites in one of the EWAS; each zoomed dot colour shows a unique CpG site specific locus in all 4 EWASs.

**Figure 2B and C.**
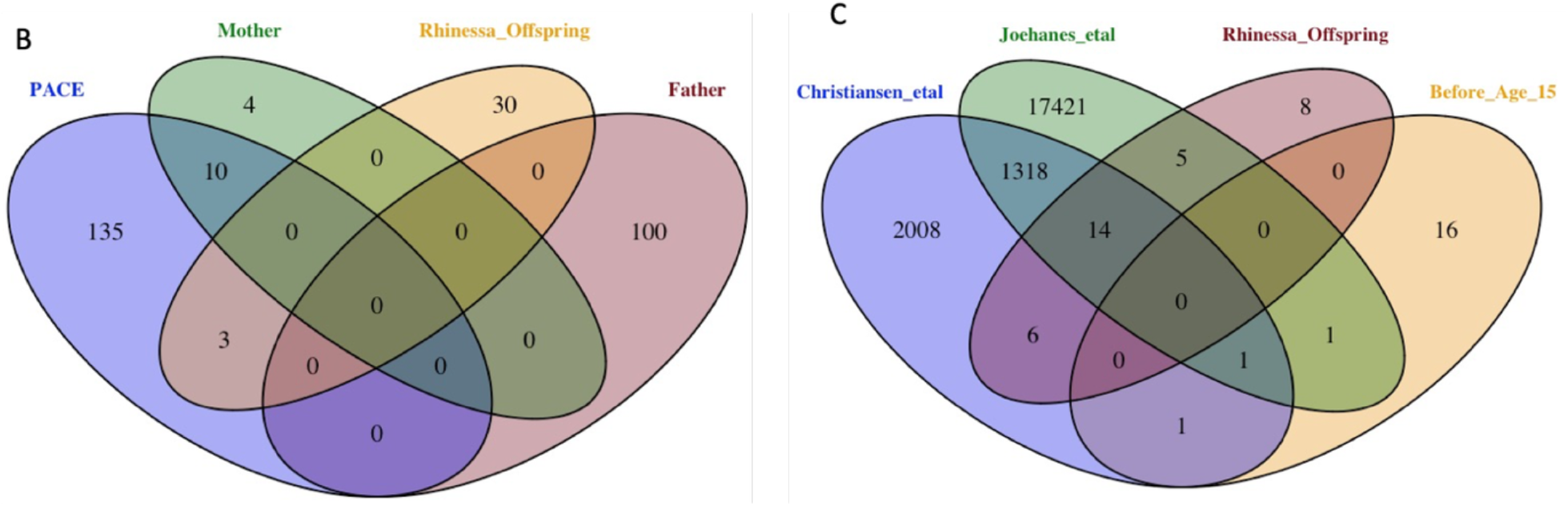
Venn diagram showing EWAS CpG top hits for personal smoking, mother’s smoking (FDR<0.005), father’s any preconception smoking (top 100 dmCpGs), and father’s pubertal smoking starting before age 15 (FDR<0.05) in the RHINESSA cohort, which are shared with top hits from meta-analysis of **B:** mother smoking (blue oval) as reported by Joubert et al 2016, and **C:** personal cigarette smoking signature as reported by Christiansen et al 2021 (blue) and by Joehanes et al 2016 (green).

Comparing our EWAS results with findings from previous studies showed that 10 of the dmCpGs we identified as related to maternal smoking, and 20 (14+6) and 19 (14+5) of the dmCpGs identified as related to personal smoking, were present in the relevant meta-analyses^16,21,22^ (Figures 2B and 2C). However, when we compared our top 100 dmCpGs for father’s any preconception smoking onset EWAS with mother’s smoking, there was no evidence for shared CpGs (Figure 2B). For father’s pubertal smoking, only two CpG sites (cg11380624 (*DNAJC14*), cg20728490 (*DNTT*)) were shared with analyses of personal smoking by Joehanes et al.^21^ and two sites (cg12053348 (intergenic), cg20728490 (*DNTT*)) with Christiansen et al. ^16^, while 16 CpG sites were unique (Figure 2C).

### Enrichment of dmCpGs for related traits

We investigated whether the significant dmCpGs associated with father’s preconception smoking onset overlapped with other traits, using the repository of published EWAS literature in the EWAS atlas. The top 23 dmCpG sites for father’s any preconception smoking (those with p-value ≤9.86 × 10^−06^, distinctly lower than the following sites) were enriched for traits that include Immunoglobulin E (IgE) level, muscle hypertrophy, maternal smoking, and birthweight (Figure 3A). dmCpGs (FDR<0.05) associated with father’s pubertal smoking were enriched for traits such as autoimmune diseases, atopy, smoking, and puberty (Figure 3B). For comparison, maternal and personal smoking dmCpGs were enriched for shared traits including aging, birthweight, cognitive function, lung function, smoking and type 2 diabetes and cancers – whereas IgE level and atopy were specifically enriched in paternal smoking (Figure 3C and 3D).

**Figure 3:**
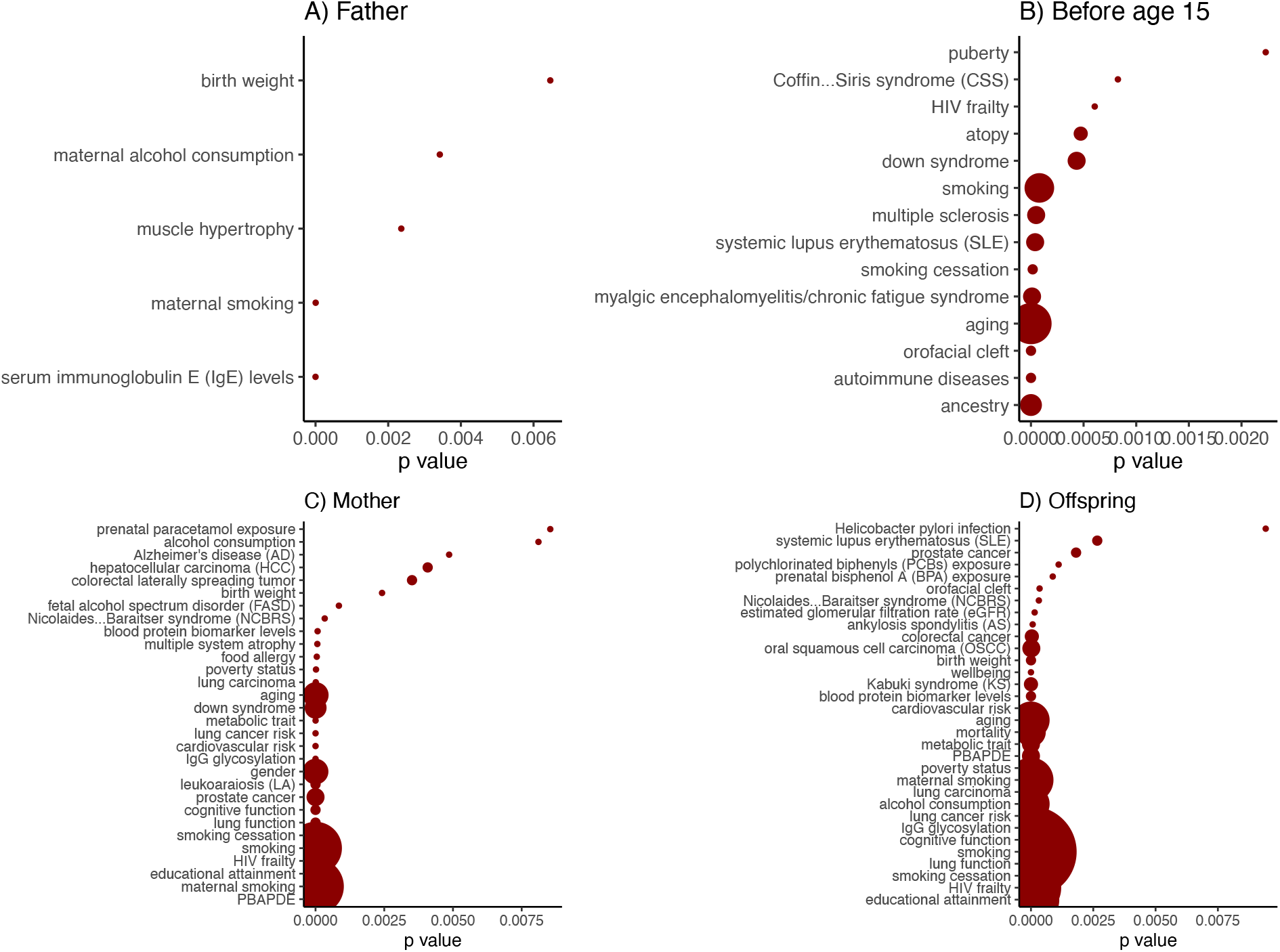
Traits associated with the CpG sites that in EWAS were identified to be differentially methylated according to **A:** father’s any preconception smoking, **B:** father’s pubertal smoking starting before age 15, **C:** Mother’s smoking and **D:** personal smoking *PPBAPDE: perinatal polychlorinated biphenyls and polychlorinated dibenzofurans exposure

### Role of dmCpGs for father’s pubertal smoking (smoking initiation < 15 years)

Given the stronger effects of father’s pubertal smoking we further explored the biological relevance of these findings.

### Transcription factor enrichment

We interrogated eFORGE TF for transcription factor enrichment in CD4^+^ cells to determine the regulatory role of our 19 significant dmCpGs (FDR<0.05) related to father’s pubertal smoking. We found significant enrichment of 27 transcription factor binding sites that overlapped with 7 of the dmCpGs (q-value<0.05) identified in our EWAS study (Supplementary Table E6).

### EWAS atlas lookup

Of the 19 dmCpGs associated with father’s pubertal smoking identified in our analysis, 11 were present in the EWAS atlas and correlated with gene expression in a variety of tissues in the EWAS atlas (Figure 4A) and overlapped with promoters (Figure 4B) (FDR <0.05). These were significantly associated with 9 other traits, including atopy and fractional exhaled nitric oxide (cg23021329), smoking (cg20728490; cg16730908), BMI (cg03516318), Acute Lymphoblastic Leukemia (cg2240207), cancer (cg11380624), and Crohn’s disease (cg10981514), (Supplementary Table E7).

**Figure 4:**
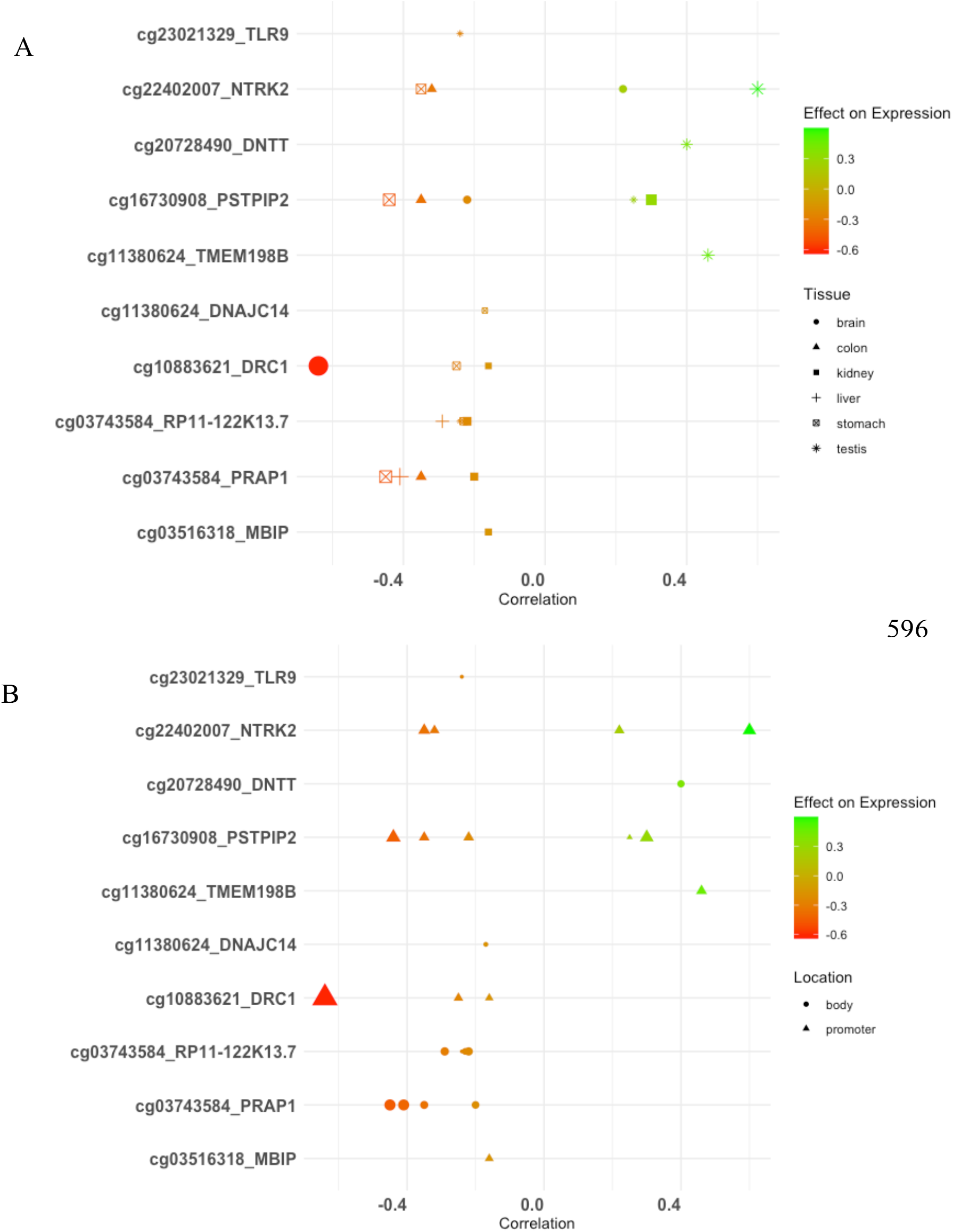
Methylation effects on gene expression regulation across different tissue types for the CpG sites differently methylated according to father’s pubertal smoking starting before age 15 years (FDR < 0.05). [Accessed on 20 June 2021]. Size of point represents -log10 p-value, colour scale shows CpG site correlation with expression; red to green represents increasing expression. In A) shape shows the tissue type, in B) shape shows genomic feature location.

### Differentially methylated region (DMR) analysis

No DMRs were significantly associated with father’s any preconception smoking using either DMRcate or dmrff. There were suggestive hits for father’s pubertal smoking, such as DNTT at FDR= 0.084. All DMRs are listed in supplementary Table E8.

### Pathway enrichment

To gain further insight into the functional roles of the dmCpGs, we used 14 genes that were mapped to dmCpGs associated with father’s pubertal smoking to generate a protein-protein interaction network from the String database. The top 20 protein interactors were included with high confidence score cutoff 0.7 from protein-protein interaction data sources including experimentally validated protein physical complexes, curated databases and co-expressions. The network indicated that immune response related genes *TLR9, CSF1R, NTRK2, PSTPIP2, PTPN11* and *IL34* were well connected (Figure 5A) (p-value <1.0×10^−16^). The molecular function enrichment analysis showed enrichment for gene expression, inflammatory response, innate immunity, and cytokine binding (Figure 5B). We also assessed enrichment of GO terms using gometh. The most significantly enriched biological process terms (FDR<0.05) include: Inactivation of MAPK activity involved in osmosensory signaling pathway (GO:0000173), negative regulation of interleukin-6 production (GO:0032715), regulation of mast cell chemotaxis (GO:0060753), regulation of neutrophil migration (GO:1902622) and insulin processing (GO:0030070) (Supplementary Table E9).

**Figure 5.**
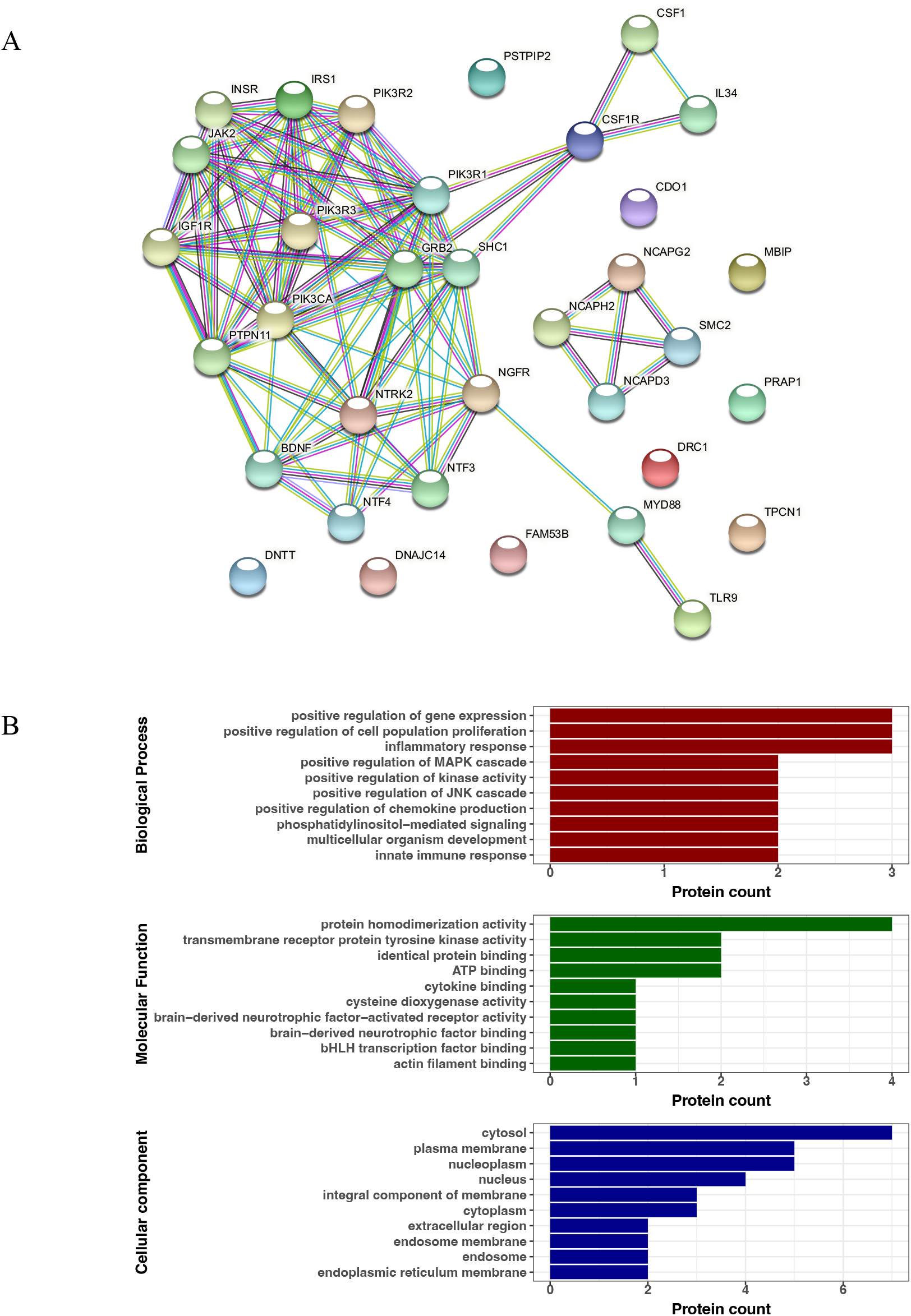
**A and B**. Interactome of dmCpGs associated with father’s pubertal smoking starting before age 15 (FDR< 0.05). **A:** Network with high confidence score 0.7 and 20 top interactors. The interaction line colour shows dataset source: Red = experimentally determined, cyan = curated database, yellow-green = text mining. **B:** Functional enrichment for gene expression regulation, inflammatory response and innate immunity.

### Replication of DNA methylation signatures associated with father’s preconception smoking

The replication cohort in ALSPAC included 542 participants (female=280, male=262), of whom 86 had a father who started to smoke before the age of 15 and 456 had never-smoking fathers. There was no overlap of dmCpG sites significantly associated with father’s smoking before age 15 between the two cohorts (FDR<0.05). However, of the 19 significant dmCpGs identified as related to father’s pubertal smoking in RHINESSA, 11 showed nominal replication in ALSPAC (p< 0.05) with similar direction. The correlation of effects between studies is R=0.49. The binomial sign test showed the association to be significant at p<0.05. Expanding the comparison to the top 100 dmCpGs in RHINESSA, the correlation of effects between studies, R = 0.54, p-value = 3.04×10^−05^.

### Sensitivity analyses

In order detect whether the associations identified were influenced by social class, we carried out regression analysis between paternal smoking associated dmCpGs as outcome and father’s education as exposure. No association was found.

In order to see the effect of CpGs changing with age, we compared known aging-related CpG markers identified from Rhinessa EWAS and paternal smoking dmCpGs with offspring age. There was only weak correlation between paternal smoking dmCpGs and offspring age (maximum R =|0.2|, with 9 CpGs R = 0). In contrast, the age-related CpG markers showed a strong correlation with age (R >=|0.6|) (Supplementary Figure 1).

In order to determine whether paternal smoking dmCpGs were associated with offspring outcomes we ran logistic and linear regression on ever-asthma, ever-wheezing, weight and BMI. Some dmCpG sites showed association with ever-asthma (cg22402007: NTRK2), ever-wheezing (cg11380624: DNAJC14, cg10981514: TPCN1), weight (cg12053348, cg03380960: FAM53B, cg22402007: NTRK2^23^) and BMI (cg03380960: FAM53B, cg12053348, cg22402007: NTRK2) at P<0.05 as shown in (Supplementary Table E10). The study power is shown in Supplementary Table E11.

## Discussion

To our knowledge, this is the first human study to investigate the potential epigenetic mechanisms behind the impact of father’s pubertal smoking on offspring. In this epigenome-wide association study, using data from two generations of study participants, we found differentially methylated CpG sites in offspring associated with father’s preconception smoking. Signatures related to father pubertal smoking (smoking initiation before age 15) were much more pronounced than smoking starting at any time preconception. Sixteen of our identified dmCpGs have not previously been reported to be associated with personal or maternal smoking. We suggest these new smoking-associated methylation biomarkers may be specific to smoking exposure of future fathers in early puberty. Several top dmCpGs were enriched for promoter regions and overlapped with significant transcription factor sites that correlated with gene expression in a variety of tissues. Besides unique sites identified for father’s preconception smoking onset, our study confirms previously reported DNA methylation sites associated with personal and mother smoking, demonstrating the validity of our cohort and analytical methods. The genes to which dmCpGs map are related to regulation of innate immunity and inflammatory responses.

For father’s any preconception smoking, we found two novel CpG sites that were not previously linked with any previously investigated smoking phenotype. PRR5 (mapped with cg008870527) is a component of the (mTOR) complex 2 which is upstream of major pathways known to have a crucial role in metabolic regulation and is suggested to play a role in obesity and the pathogenesis of insulin resistance^24^. CENPP (mapped with cg08541349), has been associated with lung function, leucocyte count, BMI and type II hypersensitivity reaction in GWAS studies^25^. In the male EWAS analysis, gene KCNJ1 is known to be associated with vital capacity and linked with obesity. A population-based study of Hispanic children has shown association of GRAMD4 with IgE levels (relevant to asthma pathogenesis)^26^. TRIM2 is linked with low density lipoprotein measurement and total cholesterol, while MYADML2 is linked with vital capacity and BMI-adjusted waist-hip ratio. Of the female EWAS hits, LEPROTL1 has a role in lung function (FEV1/FVC ratio) and several cancers, and a regulatory effect on growth hormone action and glucose homeostasis^27^.

For father’s pubertal smoking, two of our 19 significant CpG sites, have previously been associated with personal smoking (cg20728490 in *DNTT* and cg16730908 in *PSTPIP2*), and they map to genes with important roles in innate immune responses to infections^28,29^. Upregulation of *PSTPIP2* has also been linked to neutrophilic airway inflammation and non-allergic asthma. When exploring the biological impact of other genes mapped to the dmCpGs uniquely associated with father’s pubertal smoking, several were related to genes associated with innate immunity, allergic diseases, and asthma development, such as *TLR9, CSF1R, DNAJ14, NTRK2* and *TPCN1*^28–33^. We also identified CpGs and genes with links to obesity (*NTRK2, PSTPIP2, MBIP*)^25,35^, and glucose and fat metabolism (*IRS1*). The differentially methylated CpGs were mainly located in open-sea genomic features, and enriched for promoter regions, CpG island and gene bodies. These findings suggest that the identified DNA methylation differences, even though of relatively small magnitude, have functional implications in terms of a regulatory role in specific gene expression. Pathway analysis and molecular function enrichment further found interconnection of immune response related genes, and enrichment for inflammatory response, innate immunity, and cytokine binding. When seeking replication of results in an independent sample in the ALSPAC, although no dmpCpGs overlapped in the two population cohorts, results showed that effect estimates associated with fathers’ preconception smoking were moderately correlated and with concordant directional effects.

Several mechanistic reports have demonstrated that the toxicogenic components in cigarette smoke impact on epigenetic germline inheritance and affect the offspring’s metabolic health^36^. However, given this is the first study that investigated DNA methylation signatures in young and adult offspring in relation to a timing-specific exposure on father’s smoking, there is limited published literature that is directly comparable to our findings. In a pilot study, we previously observed differentially methylated regions associated with father’s ever smoking, among which annotated genes were related to innate and adaptive immunity and fatty acid synthesis^18^. Preconception paternal smoking has been shown to alter sperm DNA methylation^37^, and independently increase asthma risk and reduce lung function in the offspring ^9^, especially if the smoking started before age 15^7,9^. The observed association between the dmCpG sites related to father’s early onset smoking, and offspring asthma, wheezing and weight, suggests that epigenetic changes may lie on the casual pathway between paternal smoke exposures and offspring health outcomes.

Strikingly, the dmCpG sites we identified as related to fathers’ preconception smoking (any preconception smoking as well as pubertal smoking), were quite unique and not the same as those previously reported or found in our data to be associated with mothers’ or personal smoking. Reassuringly, our EWAS of mother’s smoking and personal smoking, identified several of the dmCpG sites related to these exposures in other cohorts.

Available data for appropriate replication of our results is a major challenge. We found moderate correlation between RHINESSA and ALSPAC EWAS for paternal smoking before 15 years. Although the replication analysis found effect estimates to have concordant directions in several of the dmCpGs, we did not identify overlapping significant dmCpGs associated with fathers’ preconception smoking in the replication cohort. The low sample size in both cohorts for paternal smoking before 15 might contribute to the lack of shared genome-wide significance. Even within the same population, using different platforms can cause difficulties with replication^38^. The similarity in the direction of association suggests a potential biological effect of early pre-puberty father’s smoking, but further research is warranted in order to verify our novel results.

Although we accounted for personal and mother’s smoking exposure in the analysis, we cannot disregard potential residual confounding related to maternal and personal smoking. Further, our analyses cannot fully disentangle effects of father’s early onset smoking from effects of subsequent accumulating second hand smoke exposure. However, epidemiological analyses of various measures of father’s smoking as related to offspring phenotype in over 20.000 father-offspring pairs found that effects of any other aspect of father’s smoking was negligible as compared to that of starting smoking early^7^. We did not control for genetic variations at single nucleotide polymorphisms and cannot rule out that the differentially methylated CpG sites are affected by, or interact with, GWAS-associated genetic variants. However, a recent analysis of our study cohorts using highly advanced statistical probabilistic simulations demonstrated that unmeasured confounding had a limited impact on the effects of father’s preconception smoking on offspring asthma^8^. This suggests that the identified dmCpGs associated with father’s preconception smoking, most likely are not driven by unmeasured confounding -by genetic factors or by lifestyle-related or environmental factors.

Self-reporting of smoking is another limitation of our study. However, based on validation studies there is an overall consensus that self-report provides a valid and reliable tool for assessing smoking behaviour in cohort studies. Furthermore, it is likely that error in father’s reporting of smoking habits is independent of DNA methylation measured in the offspring, and that misclassification thus will have attenuated the observed results and that the underlying true results might be stronger^39,40^.

We suggest that the observed association between father’s preconception smoking and offspring DNA methylation marks could be caused by transmission through germline imprint of male sperm. Supported by previous mechanistic and epidemiological findings we also speculate that our novel results reflect that early adolescence may constitute a period of particular vulnerability for smoking exposure to modify the offspring’s epigenome. A recent study demonstrated that preconception paternal cigarette exposure in mice from the onset of puberty until 2 days prior to mating modified the expression of miRNAs in spermatozoa and influenced the body weight of F1 progeny in early life^41^. As prepubertal years as well as the onset of puberty represents periods of epigenetic reprogramming events^42^, we suggest early adolescence may be a critical time for tobacco-related exposures to interfere with germline epigenetic patterns. This is, however, most challenging to study in humans and multiple scientific approaches are needed to elucidate the molecular mechanisms underlying the current findings as well as previous epidemiological results.

## Conclusion

We have identified dmCpG sites in offspring associated with father’s onset of smoking before conception, with most pronounced effects when the father started to smoke already in early puberty (before the age of 15). The pattern differed from those of maternal smoking in pregnancy and of personal smoking, and we suggest these may be unique methylation signatures specific to father’s early adolescent smoking. The genes to which the identified dmCpGs map, are related to asthma, IgE and regulation of innate immunity and inflammatory responses. Our study provide evidence for an epigenetic mechanism underlying the epidemiological findings of high risk of asthma, obesity and low lung function following father’s early adolescent smoking. The functional links of hypermethylated genes suggest that particularly father’s pubertal smoking can have cross-generational effects impacting on the long-term health in offspring. Smoking interventions in early adolescence may have implications for better public health, and potential benefits, not only for the exposed, but also for future offspring.

