## Supplementary Table for "Fathers’ preconception smoking and offspring DNA methylation: A two generation study"

Negusse T Kitaba[1*], Gerd Toril Mørkve Knudsen[2,3*], Ane Johannessen[4,] Faisal I. Rezwan[5], Andrei Malinovschi[6], Anna Oudin[7], Bryndis Benediktsdottir[8,9], David Martino[10], Francisco Javier Callejas González[11], Leopoldo Palacios Gómez[12], Mathias Holm[13], Nils Oskar Jõgi[2,3], Shyamali C Dharmage[14], Svein Magne Skulstad[3], Sarah H Watkins[15], Matthew Suderman[15], Francisco Gómez-Real[2,16], Vivi Schlünssen[17,18], Cecilie Svanes[3,4]#, John W. Holloway[1,19]#.
*Joint first authors
#Joint senior authors
[1]Human Development and Health, Faculty of Medicine, University of Southampton, Southampton, UK
[2]Department of Clinical Sciences, University of Bergen, Norway
[3]Department of Occupational Medicine, Haukeland University Hospital, Bergen, Norway
[4]Centre for International Health, Department of Global Public Health and Primary Care, University of Bergen, Norway
[5] Department of Computer Science, Aberystwyth University, Aberystwyth, UK
[6]Department of Medical Sciences: Clinical Physiology, Uppsala University, Sweden
[7]Section of Sustainable Health, Department of Public Health and Clinical Medicine, Umeå University, Sweden
[8]Department of Allergy, Respiratory Medicine and Sleep, Landspitali University Hospital, Reykjavik, Iceland
[9]Faculty of Medicine, University of Iceland, Iceland
[10]Wal-yan Respiratory Research Centre, Telethon Kids Institute, Perth, Australia
[11]Department of Pulmonology, Albacete University Hospital Complex, Albacete, Spain
[12]El Torrejón Health Centre, Andalusian Health Service, Huelva, Spain
[13]Occupational and Environmental Medicine, School of Public Health and Community Medicine, Institute of Medicine, University of Gothenburg, Sweden
[14]Centre for Epidemiology and Biostatistics, Melbourne School of Population and Global Health, University of Melbourne, Melbourne, Australia
[15]University of Bristol, MRC Integrative Epidemiology Unit, Population Health Sciences, Bristol Medical School, Bristol, United Kingdom
[16]Department of Gynaecology and Obstetrics, Haukeland University Hospital, Bergen, Norway
[17]Department of Public Health, Work, Environment and Health, Danish Ramazzini Centre, Aarhus University Denmark, Denmark
[18]National Research Center for the Working Environment, Copenhagen, Denmark
[19]NIHR Southampton Biomedical Research Center, University Hospitals Southampton, Southampton, UK

**Supplementary Methods**

*Study design and data*

Offspring were participants in the RHINESSA study (www.rhinessa.net). Parent data, including detailed information on smoking habits, were retrieved from the population-based European Community Respiratory Health Survey (ECRHS, www.ecrhs.org) and/or the Respiratory Health in Northern Europe study (RHINE, www.rhine.nu). Medical research committees in each study centre approved the studies, and each participant gave written consent. Father's smoking and age of starting/quitting was reported in interviews/questionnaires, and related to offspring's birth year, to define categories: never smoked (N=547), any preconception smoking (N=328), preconception smoking with onset <15 years (pubertal smoking) (N=64) (cut point based on mean age of voice break 14.5 years, first nocturnal seminal emission 14.8 years). Personal smoking was classified as current, ex- or never smoking. Maternal smoking was defined by offspring's report on mothers' smoking during their childhood/pregnancy.

*Methylation profiling and processing*

DNAm in offspring was measured in DNA extracted from peripheral blood, using a simple salting out procedure[1]. Bisulfite-conversion was undertaken using EZ 96-DNA methylation kits (Zymo Research, Irvine, CA, USA) at the Oxford Genomics Centre (Oxford, UK) and methylation assessed using Illumina Infinium MethylationEPIC Beadchip arrays (Illumina, Inc. CA, USA) with samples randomly distributed on microarrays to control against batch effects.

Data analysis was undertaken using Statistical Computing Program R, version 3.6.1[2]. Methylation intensity files were processed and quality was assessed using minfi[3] and Mefil[4]. Methylation distribution for outliers were assessed using density and multidimensional scaling plots, methylated vs unmethylated ratio plot, sex mismatch and sex outliers, control probes and bisulphite conversion efficiency.

Normalization was carried out using BMIQ, which adjusts intra-sample the beta-values of type 2 design and type 1 probes[5]. To remove technical variation detected by champ.SVD function within the CHAMP package[6], ComBat from SVA was applied on plate and slides for batch effect correction[7]. Probes were excluded from analysis using the following criteria: detection p-value above 0.01 (n=24566 probes), probes associated with SNP, probes with a beadcount <3 in at least 5% of samples (n=1437), multiple locations, non-cg probes (n=2624), probes on the X or Y chromosomes (n= 16556) and cross-reactive probes on the EPIC array (n=43000)[8]. Cell-type proportion was estimated with EpiDISH (epigenetics Dissection of Intra-Sample Heterogeneity) [9]. Following processing, 726,661 CpGs were retained for analysis.

*Statistical analysis*

We ran two EWAS on preconception father's smoking as exposure (any preconception smoking, and prepuberty smoking) with DNA methylation as outcome. To identify differentially-methylated Cytosine-phosphate-Guanine (CpG) sites (dmCPG), robust multiple linear regression models were applied on beta values using limma[10] adjusting for offspring's sex, age, personal and mother's smoking, and cell-type proportions (B-cells, Natural killer cells, CD4 T-cells, CD8 T-cells, Monocyte, Neutrophils) at significance level of false discovery rate (FDR)[11] corrected p-value<0.05. Eosinophils were not included due to a very low estimate. Manhattan plots were generated using qqman[12] and circos plot with CMplot R package[13]. Inflation from systematic biases was adjusted using BACON[14]. Differentially methylated regions were detected using dmrff[15]. Transcription factor binding site prediction was performed using eFORGE TF[16]. Gene-disease association was identified using open target[17]. Identified dmCpGs were compared against EWAS atlas for association with known biological traits[18]. To gain biological insight regarding the dmCpGs mapped to genes, gene interactors were identified using String[19] and enrichment was performed using UniprotR[20] and gometh[21].

We compared our EWAS results with findings from meta-analysis of EPIC DNA methylation associated with personal smoking from four population-based cohorts[22], personal smoking-methylation effects from 16 cohorts using 450K arrays[23]; and the Pregnancy and Childhood Epigenetics Consortium (PACE) meta-analysis of mother smoking on offspring cordblood methylation[24].

**ALSPAC Cohort description**

ALSPAC is a pre-birth cohort designed to determine the environmental and genetic factors that are associated with health and development of the study offspring (1-3). Ethical approval for the study was obtained from the ALSPAC Ethics and Law Committee and the Local Research Ethics Committees (4). Consent for biological samples has been collected in accordance with the Human Tissue Act (2004). Informed consent for the use of data collected via questionnaires and clinics was obtained from participants following the recommendations of the ALSPAC Ethics and Law Committee at the time.

Pregnant women resident in Avon, UK, with expected dates of delivery 1st April 1991 to 31st December 1992 were invited to take part in the study. The initial number of pregnancies enrolled is 14,541 (for these at least one questionnaire has been returned or a “Children in Focus” clinic had been attended by 19/07/99). Of these initial pregnancies, there was a total of 14,676 fetuses, resulting in 14,062 live births and 13,988 children who were alive at 1 year of age.

When the oldest children were approximately 7 years of age, an attempt was made to bolster the initial sample with eligible cases who had failed to join the study originally. As a result, when considering variables collected from the age of seven onwards (and potentially abstracted from obstetric notes) there are data available for more than the 14,541 pregnancies mentioned above. The number of new pregnancies not in the initial sample (known as Phase I enrolment) that are currently represented on the built files and reflecting enrolment status at the age of 24 is 913 (456, 262 and 195 recruited during Phases II, III and IV respectively), resulting in an additional 913 children being enrolled. The phases of enrolment are described in more detail in the cohort profile paper and its update (2, 3). The total sample size for analyses using any data collected after the age of seven is therefore 15,454 pregnancies, resulting in 15,589 fetuses. Of these 14,901 were alive at 1 year of age.

Please note that the study website contains details of all the data that is available through a fully searchable data dictionary and variable search tool: http://www.bristol.ac.uk/alspac/researchers/our-data/

ALSPAC fully supports Wellcome and the RCUK policies on open access. The process for obtaining access to data is described on the study website: http://www.bristol.ac.uk/alspac/researchers/data-access/. The datasets for this study will not be made publicly available, as in order to preserve confidentiality of the participants it is important that the ALSPAC access rules are taken into account. The ALSPAC study website contains details of all the data that are available through a fully searchable data dictionary: http://www.bristol.ac.uk/alspac/researchers/our-data/.

1. Boyd A, Golding J, Macleod J, Lawlor DA, Fraser A, Henderson J, Molloy L, Ness A, Ring S, Davey Smith G. Cohort Profile: The 'Children of the 90s'; the index offspring of The Avon Longitudinal Study of Parents and Children (ALSPAC). International Journal of Epidemiology 2013; 42: 111-127.
2. Fraser A, Macdonald-Wallis C, Tilling K, Boyd A, Golding J, Davey Smith G, Henderson J, Macleod J, Molloy L, Ness A, Ring S. Cohort profile: the Avon Longitudinal Study of Parents and Children: ALSPAC mothers cohort. International journal of epidemiology. 2013 Feb 1;42(1):97-110.
3. Northstone K, Lewcock M, Groom A, Boyd A, Macleod J, Timpson NJ, Wells N. The Avon Longitudinal Study of Parents and Children (ALSPAC): an updated on the enrolled sample of index children in 2019. Wellcome Open research 2019; 4:51 (https://doi.org/10.12688/wellcomeopenres.15132.1)
4. Birmingham, K. (2018). *Pioneering ethics in longitudinal studies: The early development of the ALSPAC Ethics & Law Committee*. Bristol: Policy Press

**Acknowledgements**

We are extremely grateful to all the families who took part in this study, the midwives for their help in recruiting them, and the whole ALSPAC team, which includes interviewers, computer and laboratory technicians, clerical workers, research scientists, volunteers, managers, receptionists and nurses.

**Funding of ALSPAC**

The UK Medical Research Council and Wellcome Trust (Grant ref: 217065/Z/19/Z) and the University of Bristol currently provide core support for ALSPAC. A comprehensive list of grants funding is available on the ALSPAC website

(http://www.bristol.ac.uk/alspac/external/documents/grant-acknowledgements.pdf).

Supplementary Table E1: Top dmCpG sites associated with fathers' any preconception smoking onset (N=875)

| CpG | Coefficient | Average | P- value | Adj.P | Chromosome | Location | Gene | Ref Gene |
|---|---|---|---|---|---|---|---|---|
| **cg00870527** | -0.02 | 0.50 | 3.97E-08 | 0.03 | 22 | N_Shelf | PRR5 | 5'UTR |
| **cg08541349** | -0.01 | 0.88 | 7.92E-08 | 0.03 | 9 | OpenSea | CENPP | Body |
| **cg01000965** | 0.00 | 0.94 | 3.70E-07 | 0.09 | 16 | OpenSea | NPRL3 | Body |
| **cg06115838** | -0.09 | 0.53 | 7.09E-07 | 0.10 | 17 | S_Shelf | cg06115838 | |
| **cg12819747** | 0.00 | 0.94 | 7.15E-07 | 0.10 | 12 | Island | C12orf51 | Body |
| **cg25390635** | -0.01 | 0.94 | 1.40E-06 | 0.17 | 22 | Island | GRAMD4 | TSS200 |
| **cg00061114** | -0.02 | 0.76 | 2.38E-06 | 0.19 | 2 | OpenSea | CCDC88A | Body |
| **cg19646139** | -0.01 | 0.93 | 3.12E-06 | 0.19 | 1 | S_Shelf | PRKCZ | Body |
| **cg26884359** | -0.01 | 0.84 | 3.38E-06 | 0.19 | 17 | N_Shelf | cg26884359 | |
| **cg22778120** | -0.02 | 0.54 | 3.50E-06 | 0.19 | 22 | N_Shelf | PRR5 | 5'UTR |
| **cg06354075** | -0.01 | 0.90 | 4.01E-06 | 0.19 | 11 | OpenSea | NUP98 | Body |
| **cg05458764** | -0.01 | 0.92 | 4.12E-06 | 0.19 | 5 | Island | TPPP | Body |
| **cg25817279** | -0.01 | 0.83 | 4.19E-06 | 0.19 | 2 | OpenSea | cg25817279 | |
| **cg24284924** | 0.00 | 0.95 | 4.21E-06 | 0.19 | 6 | OpenSea | cg24284924 | |
| **cg06597248** | -0.02 | 0.62 | 4.34E-06 | 0.19 | 2 | OpenSea | cg06597248 | |
| **cg09543474** | 0.00 | 0.93 | 4.47E-06 | 0.19 | 5 | OpenSea | CNOT6 | Body |
| **cg17931890** | 0.00 | 0.07 | 4.56E-06 | 0.19 | 2 | Island | cg17931890 | |
| **cg17545662** | -0.01 | 0.94 | 5.13E-06 | 0.21 | 6 | OpenSea | C6orf70 | Body |
| **cg06868293** | -0.01 | 0.92 | 6.07E-06 | 0.23 | 2 | OpenSea | SCN2A | Body |
| **cg10231096** | -0.01 | 0.92 | 6.65E-06 | 0.24 | 15 | OpenSea | TRPM1 | Body |
| **cg25710809** | -0.01 | 0.79 | 7.08E-06 | 0.24 | 8 | OpenSea | GNRH1 | TSS1500 |
| **cg03657121** | -0.01 | 0.82 | 8.70E-06 | 0.25 | 7 | OpenSea | UBE3C | Body |
| **cg09831081** | -0.01 | 0.87 | 9.86E-06 | 0.25 | 21 | N_Shore | LSS | Body |

Supplementary Table E2: Sex-stratified dmCpGs (FDR<=0.05) associated with fathers' any preconception smoking in male (N=457) and female (N=418) offspring

| Offspring sex | CpG | Coefficient | Average | P- value | Adj.P | Chr | Location | Gene | Ref Gene |
|---|---|---|---|---|---|---|---|---|---|
| Males | cg05193832 | -0.01 | 0.94 | 5.16E-08 | 0.04 | 11 | OpenSea | KCNJ1 | 5'UTR |
| | cg25390635 | -0.01 | 0.94 | 1.05E-07 | 0.04 | 22 | Island | GRAMD4 | TSS200 |
| | cg22905274 | -0.01 | 0.04 | 2.55E-07 | 0.05 | 4 | Island | TRIM2 | Body |
| | cg02518394 | -0.01 | 0.91 | 2.87E-07 | 0.05 | 17 | OpenSea | MYADML2 | TSS1500 |
| Females | cg09801901 | -0.02 | 0.71 | 3.18E-08 | 0.02 | 8 | OpenSea | LEPROTL1 | Body |

Supplementary Table E3: dmCpGs associated with father's smoking onset before age 15 years (N=304, FDR<0.05)

| CpG | Coefficient | Average | P-value | Adj.P | Chromosome | Location | Gene | Ref Gene |
|---|---|---|---|---|---|---|---|---|
| **cg23021329** | 0.01 | 0.27 | 6.07E-08 | 0.03 | 3 | S_Shore | TLR9 | Body |
| **cg20728490** | 0.03 | 0.37 | 6.30E-08 | 0.03 | 10 | OpenSea | DNTT | 5'UTR |
| **cg12053348** | 0.04 | 0.61 | 6.55E-08 | 0.03 | 3 | OpenSea | cg12053348 | |
| **cg03380960** | 0.02 | 0.48 | 1.11E-07 | 0.03 | 10 | OpenSea | FAM53B | Body |
| **cg26274304** | 0.02 | 0.36 | 1.66E-07 | 0.04 | 7 | N_Shore | NCAPG2 | 5'UTR |
| **cg16730908** | 0.02 | 0.39 | 2.08E-07 | 0.04 | 18 | S_Shore | PSTPIP2 | TSS1500 |
| **cg13904562** | 0.04 | 0.53 | 2.26E-07 | 0.04 | 12 | OpenSea | cg13904562 | |
| **cg07508217** | 0.03 | 0.69 | 2.60E-07 | 0.04 | 3 | OpenSea | cg07508217 | |
| **cg03516318** | 0.03 | 0.21 | 2.87E-07 | 0.04 | 14 | OpenSea | MBIP | TSS1500 |
| **cg10883621** | 0.02 | 0.35 | 3.00E-07 | 0.04 | 2 | Island | C2orf39 | TSS200 |
| **cg22402007** | 0.02 | 0.16 | 3.88E-07 | 0.04 | 9 | N_Shore | NTRK2 | TSS1500 |
| **cg11380624** | 0.02 | 0.27 | 4.04E-07 | 0.04 | 12 | N_Shore | DNAJC14 | 5'UTR |
| **cg15882605** | 0.02 | 0.44 | 4.60E-07 | 0.04 | 22 | OpenSea | cg15882605 | |
| **cg03818156** | 0.02 | 0.91 | 4.62E-07 | 0.04 | 19 | OpenSea | cg03818156 | |
| **cg13288863** | 0.02 | 0.79 | 5.93E-07 | 0.05 | 5 | N_Shore | CDO1 | Body |
| **cg03743584** | 0.02 | 0.30 | 6.33E-07 | 0.05 | 10 | OpenSea | PRAP1 | 1stExon |
| **cg10981514** | 0.02 | 0.42 | 6.49E-07 | 0.05 | 12 | OpenSea | TPCN1 | Body |
| **cg06600694** | 0.00 | 0.06 | 6.82E-07 | 0.05 | 2 | Island | IRS1 | TSS200 |
| **cg14700085** | 0.02 | 0.71 | 7.56E-07 | 0.05 | 5 | OpenSea | CSF1R | Body |
| **cg25406294** | 0.02 | 0.78 | 8.42E-07 | 0.05 | 14 | OpenSea | cg25406294 | |

Supplementary Table E4: dmCpGs associated with offspring's own smoking status (N=875, FDR<0.05)

| CpG | Coefficient | Average | P-value | Adj.P | Chromosome | Location | Gene | Ref Gene |
|---|---|---|---|---|---|---|---|---|
| **cg05575921** | -0.04 | 0.83 | 2.68E-54 | 1.94E-48 | 5 | N_Shore | AHRR | Body |
| **cg21566642** | -0.04 | 0.55 | 8.21E-29 | 2.97E-23 | 2 | Island | cg21566642 | |
| **cg06644428** | -0.03 | 0.12 | 5.35E-20 | 1.29E-14 | 2 | Island | cg06644428 | |
| **cg01940273** | -0.03 | 0.58 | 1.50E-16 | 2.72E-11 | 2 | Island | cg01940273 | |
| **cg17739917** | -0.03 | 0.39 | 4.85E-16 | 6.77E-11 | 17 | S_Shelf | RARA | 5'UTR |
| **cg03636183** | -0.03 | 0.63 | 5.61E-16 | 6.77E-11 | 19 | N_Shore | F2RL3 | Body |
| **cg21911711** | -0.02 | 0.84 | 7.79E-12 | 8.05E-07 | 19 | N_Shore | F2RL3 | TSS1500 |
| **cg25189904** | -0.03 | 0.38 | 6.96E-11 | 6.30E-06 | 1 | S_Shore | GNG12 | TSS1500 |
| **cg12806681** | -0.01 | 0.94 | 1.31E-10 | 1.06E-05 | 5 | N_Shore | AHRR | Body |
| **cg21161138** | -0.02 | 0.73 | 1.60E-10 | 1.16E-05 | 5 | OpenSea | AHRR | Body |
| **cg09338374** | 0.02 | 0.49 | 9.29E-10 | 6.11E-05 | 22 | S_Shelf | cg09338374 | |
| **cg24838345** | -0.02 | 0.84 | 1.44E-09 | 8.13E-05 | 8 | N_Shelf | MTSS1 | Body |
| **cg26703534** | -0.02 | 0.68 | 1.56E-09 | 8.13E-05 | 5 | S_Shelf | AHRR | Body |
| **cg14753356** | -0.022 | 0.41 | 1.57E-09 | 8.13E-05 | 6 | OpenSea | cg14753356 | |
| **cg18110140** | -0.024 | 0.44 | 1.95E-08 | 0.00094207 | 15 | OpenSea | cg18110140 | |
| **cg26718213** | 0.035 | 0.22 | 3.50E-08 | 0.00158405 | 2 | Island | SNED1 | Body |
| **cg05086879** | -0.016 | 0.83 | 6.03E-08 | 0.00256768 | 22 | OpenSea | MGAT3 | 5'UTR |
| **cg21322436** | -0.01 | 0.25 | 6.71E-08 | 0.00269941 | 7 | N_Shore | CNTNAP2 | TSS1500 |
| **cg25648203** | -0.015 | 0.80 | 9.27E-08 | 0.00352967 | 5 | OpenSea | AHRR | Body |
| **cg20832643** | -0.014 | 0.89 | 1.23E-07 | 0.00428032 | 6 | OpenSea | TRAF3IP2-AS1 | Body |
| **cg09935388** | -0.026 | 0.76 | 1.24E-07 | 0.00428032 | 1 | Island | GFI1 | Body |
| **cg18096787** | 0.016 | 0.59 | 2.44E-07 | 0.00803438 | 21 | OpenSea | cg18096787 | |
| **cg24090911** | -0.018 | 0.75 | 3.31E-07 | 0.01041909 | 5 | OpenSea | AHRR | Body |
| **cg19859270** | -0.006 | 0.95 | 4.17E-07 | 0.01258376 | 3 | OpenSea | GPR15 | 1stExon |
| **cg25159376** | -0.002 | 0.021 | 5.53E-07 | 0.01600192 | 14 | Island | KLHDC1 | TSS200 |
| **cg17025708** | -0.006 | 0.92 | 6.49E-07 | 0.01783236 | 10 | OpenSea | VTI1A | Body |
| **cg02978227** | -0.006 | 0.94 | 6.65E-07 | 0.01783236 | 3 | OpenSea | cg02978227 | |
| **cg26707709** | 0.017 | 0.11 | 1.28E-06 | 0.03257775 | 2 | Island | SNED1 | Body |
| **cg23577033** | -0.005 | 0.932 | 1.31E-06 | 0.03257775 | 10 | S_Shelf | cg23577033 | |

| | | | | | | | | |
|---|---|---|---|---|---|---|---|---|
| **cg25401612** | -0.03 | 0.72 | 2.02E-06 | 0.04794569 | 12 | OpenSea | cg25401612 | |
| **cg13525276** | 0.02 | 0.27 | 2.05E-06 | 0.04794569 | 14 | OpenSea | TSHR | Body |
| **cg17260354** | 0.014 | 0.45 | 2.13E-06 | 0.04819969 | 17 | N_Shore | CDK3 | 3'UTR |
| **cg06117824** | -0.004 | 0.96 | 2.28E-06 | 0.04990539 | 1 | S_Shelf | TMEM51 | Body |
| **cg25949550** | -0.006 | 0.086 | 2.43E-06 | 0.05178064 | 7 | S_Shore | CNTNAP2 | Body |
| **cg15342087** | -0.006 | 0.914 | 2.74E-06 | 0.05448434 | 6 | OpenSea | | |
| **cg22635676** | 0.046 | 0.24 | 2.79E-06 | 0.05448434 | 2 | Island | SNED1 | Body |
| **cg21747070** | -0.01 | 0.52 | 2.79E-06 | 0.05448434 | 5 | N_Shore | | |

Supplementary Table E5: dmCpGs associated with mothers' smoking during offspring's childhood (N=875, FDR<0.05)

| CpG | Coefficient | Average | P-value | Adj.P | Chromosome | Location | Gene | Ref Gene |
|---|---|---|---|---|---|---|---|---|
| **cg19089201** | 0.03 | 0.71746013 | 1.47E-10 | 0.0001 | 7 | Island | MYO1G | 3'UTR |
| **cg12803068** | 0.06 | 0.67728595 | 3.31E-10 | 0.0001 | 7 | S_Shore | MYO1G | Body |
| **cg05549655** | 0.02 | 0.18071069 | 6.91E-09 | 0.0015 | 15 | Island | CYP1A1 | TSS1500 |
| **cg14179389** | -0.03 | 0.26117365 | 8.48E-09 | 0.0015 | 1 | Island | GFI1 | Body |
| **cg04180046** | 0.03 | 0.45264404 | 1.73E-08 | 0.0025 | 7 | Island | MYO1G | Body |
| **cg25949550** | -0.01 | 0.08628854 | 2.10E-08 | 0.0025 | 7 | S_Shore | CNTNAP2 | Body |
| **cg13305373** | 0.01 | 0.91366976 | 5.71E-08 | 0.0059 | 17 | OpenSea | RGS9 | Body |
| **cg22549041** | 0.03 | 0.32680072 | 1.63E-07 | 0.0147 | 15 | Island | CYP1A1 | TSS1500 |
| **cg05009104** | 0.03 | 0.69281994 | 2.53E-07 | 0.0180 | 7 | S_Shore | MYO1G | Body |
| **cg11924019** | 0.02 | 0.3723796 | 2.63E-07 | 0.0180 | 15 | Island | CYP1A1 | TSS1500 |
| **cg12101586** | 0.03 | 0.47187709 | 2.74E-07 | 0.0180 | 15 | Island | CYP1A1 | TSS1500 |
| **cg05777089** | 0.01 | 0.08624407 | 6.30E-07 | 0.0380 | 1 | OpenSea | BRINP3 | 5'UTR |
| **cg13570656** | 0.03 | 0.38212374 | 6.84E-07 | 0.0381 | 15 | Island | CYP1A1 | TSS1500 |
| **cg04785284** | 0.01 | 0.9528626 | 8.18E-07 | 0.0423 | 16 | OpenSea | cg04785284 | |

Supplementary Table E6: Transcription factor Enrichment (q-value<0.05) for dmCpGs (FDR<0.05) associated with fathers' smoking onset before age 15 years

| TF | Database | p-value | q-value | CpGs | Gene |
|---|---|---|---|---|---|
| **V_GZF1_01** | TRANSFAC | 7.09E-06 | 0.00042543 | cg22402007 | NTRK2 |
| **HSF1_HSF_2** | Taipale/SELEX | 7.70E-06 | 0.00042543 | cg12053348 | N/A |
| **HSF4_HSF_1** | Taipale/SELEX | 8.88E-06 | 0.00042543 | cg12053348 | N/A |
| **HSF1_HSF_1** | Taipale/SELEX | 1.03E-05 | 0.00042543 | cg12053348 | N/A |
| **MA0090.1-TEAD1** | JASPAR | 4.06E-05 | 0.00091734 | cg10981514 | TPCN1 |
| **V_TEF_01** | TRANSFAC | 4.06E-05 | 0.00091734 | cg10981514 | TPCN1 |
| **V_ELK1_03** | TRANSFAC | 4.60E-05 | 0.00091734 | cg10981514 | TPCN1 |
| **V_TAL1BETAITF2_01** | TRANSFAC | 4.67E-05 | 0.00091734 | cg16730908 | PSTPIP2 |
| **V_TAL1BETAE47_01** | TRANSFAC | 4.98E-05 | 0.00091734 | cg16730908 | PSTPIP2 |
| **V_ELK1_04** | TRANSFAC | 8.53E-05 | 0.00141511 | cg10981514 | TPCN1 |
| **V_HSF1_Q6** | TRANSFAC | 0.00014091 | 0.00212498 | cg12053348 | N/A |
| **Ets1** | UniProbe | 0.00015508 | 0.00214386 | cg10981514 | TPCN1 |
| **MA0028.1-ELK1** | JASPAR | 0.00026585 | 0.00330968 | cg10981514 | TPCN1 |
| **V_ETS_Q4** | TRANSFAC | 0.00027932 | 0.00330968 | cg15882605 | N/A |
| **Tcfe2a_secondary** | UniProbe | 0.00045637 | 0.00504704 | cg16730908 | PSTPIP2 |
| **ETV6_ETS_1** | Taipale/SELEX | 0.00071753 | 0.00743932 | cg10981514 | TPCN1 |
| **MA0079.2-SP1** | JASPAR | 0.00083946 | 0.00758437 | cg11380624/ cg06600694 | DNAJC14 / IRS1 |
| **V_ZBP89_Q4** | TRANSFAC | 0.00086685 | 0.00758437 | cg11380624 | DNAJC14 |
| **V_GADP_01** | TRANSFAC | 0.00086868 | 0.00758437 | cg15882605 | N/A |
| **V_GABP_B** | TRANSFAC | 0.00114402 | 0.00948899 | cg15882605 | N/A |
| **V_SP1_Q2_01** | TRANSFAC | 0.00197707 | 0.0156178 | cg15882605 | N/A |
| **V_SP4_Q5** | TRANSFAC | 0.00257797 | 0.0194388 | cg11380624/ cg06600694 | DNAJC14/ IRS1 |
| **Zfp740_primary** | UniProbe | 0.0036371 | 0.0262327 | cg06600694 | IRS1 |

Supplementary Table E7: EWAS Atlas lookup for replication (accessed on 20 June 2021) for top dmCpGs.

| CpG | Gene | Traits |
|---|---|---|
| **cg00870527** | PRR5 | Gulf War Illness and Papuan ancestry proportions |
| **cg08541349** | CENPP | No report |
| **cg01000965** | NPRL3 | No report |
| **cg05193832** | KCNJ1 | No report |
| **cg25390635** | GRAMD4 | serum immunoglobulin E (IgE) levels |
| **cg22905274** | TRIM2 | No report |
| **cg02518394** | MYADML2 | autism spectrum disorders (ASD) |
| **cg09801901** | LEPROTL1 | No report |
| **cg23021329** | TLR9 | atopy and fractional exhaled nitric oxide |
| **cg20728490** | DNTT | Smoking, aging |
| **cg12053348** | N/A | Aging |
| **cg03380960** | FAM53 | Aging |
| **cg26274304** | NCAPG2 | No report |
| **cg16730908** | PSTPIP2 | Smoking |
| **cg13904562** | N/A | No report |
| **cg07508217** | N/A | No report |
| **cg03516318** | MBIP | BMI |
| **cg10883621** | C2orf39 | B Acute Lymphoblastic Leukaemia with t(1;19)(q23;p13.3); E2A-PBX1 (TCF3-PBX1) |
| **cg22402007** | NTRK2 | B Acute Lymphoblastic Leukaemia with t(12;21)(p13.2;q22.1); ETV6-RUNX1 |
| **cg11380624** | DNAJC14 | Cancer |
| **cg15882605** | N/A | No report |
| **cg03818156** | N/A | No report |
| **cg13288863** | CDO1 | No report |
| **cg03743584** | PRAP1 | AD |
| **cg10981514** | TPCN1 | Crohn's disease |
| **cg06600694** | IRS1 | No report |
| **cg14700085** | CSF1R | No report |

Supplementary Table E8: Differentially Methylated Regions for father smoking onset before age 15 years.

| Region | Site | Chr | Position | Gene | Estimate | P-value | dmr.start | dmr.end | dmr.z | dmr P.adj. |
|---|---|---|---|---|---|---|---|---|---|---|
| **20599** | 635922 | 10 | 98064175 | DNTT | 0.03171911 | 1.04E-07 | 98064175 | 98064175 | 5.32642756 | 0.0841472 |
| **43571** | 649032 | 3 | 53461190 | | 0.02527496 | 1.05E-07 | 53461190 | 53461190 | 5.32370777 | 0.08541576 |
| **6282** | 253453 | 10 | 126390659 | FAM53B | 0.00914814 | 0.00847902 | 126390317 | 126390659 | 5.25221053 | 0.12625419 |
| **6282** | 504848 | 10 | 126390317 | FAM53B | 0.0063052 | 0.02895291 | 126390317 | 126390659 | 5.25221053 | 0.12625419 |
| **24736** | 585899 | 12 | 56222881 | DNAJC14 | 0.02359829 | 2.28E-07 | 56222881 | 56222881 | 5.17393919 | 0.19255909 |
| **27284** | 207465 | 14 | 61070289 | | 0.02108314 | 2.39E-07 | 61070289 | 61070289 | 5.16542124 | 0.20153858 |
| **2688** | 141869 | [illegible] | 48690974 | | 0.04041497 | 4.05E-07 | 48690974 | 48691036 | 5.15848642 | 0.20914664 |
| **2688** | 681241 | 12 | 48691036 | | 0.02183902 | 7.20E-06 | 48690974 | 48691036 | 5.15848642 | 0.20914664 |
| **34971** | 273463 | 19 | 3073159 | | 0.01729797 | 3.68E-07 | 3073159 | 3073159 | 5.08022606 | 0.31670631 |
| **25389** | 341664 | 12 | 113678406 | TPCN1 | 0.02375583 | 4.26E-07 | 113678406 | 113678406 | 5.05092116 | 0.36937942 |
| **3443** | 273124 | 3 | 72536534 | | 0.03521499 | 4.93E-07 | 72536534 | 72536534 | 5.0218192 | 0.43000027 |
| **705** | 350648 | 17 | 76130139 | TMC8 | 0.02054525 | 4.64E-06 | 76129984 | 76130305 | 5.00220297 | 0.47615976 |
| **705** | 647240 | 17 | 76130305 | TMC8 | 0.01594663 | 6.66E-06 | 76129984 | 76130305 | 5.00220297 | 0.47615976 |
| **705** | 405230 | 17 | 76129984 | TMC8;TMC6 | 0.01500805 | 4.97E-05 | 76129984 | 76130305 | 5.00220297 | 0.47615976 |
| **47604** | 378684 | 5 | 115149472 | CDO1 | 0.02026539 | 5.81E-07 | 115149472 | 115149472 | 4.98898749 | 0.50991255 |
| **9368** | 247050 | 3 | 159570903 | SCHIP1 | 0.01794319 | 6.27E-07 | 159570903 | 159570903 | 4.97338472 | 0.55273525 |
| **324** | 705691 | 17 | 58499720 | C17orf64 | 0.02077587 | 1.28E-05 | 58499679 | 58499720 | 4.92454708 | 0.71035553 |
| **324** | 504219 | 17 | 58499706 | C17orf64 | 0.02006096 | 7.75E-05 | 58499679 | 58499720 | 4.92454708 | 0.71035553 |
| **324** | 30671 | 17 | 58499679 | C17orf64 | 0.01173987 | 0.00023255 | 58499679 | 58499720 | 4.92454708 | 0.71035553 |
| **324** | 36291 | 17 | 58499700 | C17orf64 | 0.0137547 | 0.00770448 | 58499679 | 58499720 | 4.92454708 | 0.71035553 |
| **7044** | 542568 | 14 | 36790242 | MBIP;MBIP | 0.02684703 | 8.04E-07 | 36790242 | 36790242 | 4.92331005 | 0.71486272 |
| **9085** | 389031 | 22 | 37562418 | | 0.02544397 | 8.98E-07 | 37562418 | 37562418 | 4.90066973 | 0.80238468 |

Supplementary Table E9: Gene Ontology Terms from gometh for dmCpGs (FDR<0.05) associated with fathers' smoking onset before age 15years. BP: biological processing; MF: molecular function; CC cellular components; N: number of genes in GO term; De: number of differentially methylated genes; P.De: P-value for over-representation of GO term

| GO_name | ONTOLOGY | TERM | N | DE | P.DE |
|---|---|---|---|---|---|
| **GO:0000173** | BP | inactivation of MAPK activity involved in osmosensory signaling pathway | 1 | 1 | 0.001 |
| **GO:0032715** | BP | negative regulation of interleukin-6 production | 35 | 2 | 0.001 |
| **GO:0003912** | MF | DNA nucleotidylexotransferase activity | 1 | 1 | 0.001 |
| **GO:0060753** | BP | regulation of mast cell chemotaxis | 1 | 1 | 0.001 |
| **GO:1902622** | BP | regulation of neutrophil migration | 1 | 1 | 0.001 |
| **GO:0048709** | BP | oligodendrocyte differentiation | 24 | 2 | 0.002 |
| **GO:0004860** | MF | protein kinase inhibitor activity | 27 | 2 | 0.002 |
| **GO:0060175** | MF | brain-derived neurotrophic factor-activated receptor activity | 1 | 1 | 0.002 |
| **GO:0036019** | CC | Endolysosome | 1 | 1 | 0.002 |
| **GO:0044147** | BP | negative regulation of development of symbiont involved in interaction with host | 1 | 1 | 0.003 |
| **GO:0052362** | BP | catabolism by host of symbiont protein | 1 | 1 | 0.003 |
| **GO:1905597** | BP | positive regulation of low-density lipoprotein particle receptor binding | 1 | 1 | 0.003 |
| **GO:1905599** | BP | positive regulation of low-density lipoprotein receptor activity | 1 | 1 | 0.003 |
| **GO:0006304** | BP | DNA modification | 2 | 1 | 0.003 |

| | | | | | |
|---|---|---|---|---|---|
| **GO:0045356** | BP | positive regulation of interferon-alpha biosynthetic process | 4 | 1 | 0.003 |
| **GO:0010810** | BP | regulation of cell-substrate adhesion | 2 | 1 | 0.003 |
| **GO:0034163** | BP | regulation of toll-like receptor 9 signaling pathway | 2 | 1 | 0.004 |
| **GO:0006043** | BP | glucosamine catabolic process | 2 | 1 | 0.004 |
| **GO:0052405** | BP | negative regulation by host of symbiont molecular function | 3 | 1 | 0.004 |
| **GO:0019471** | BP | 4-hydroxyproline metabolic process | 3 | 1 | 0.004 |
| **GO:0019834** | MF | phospholipase A2 inhibitor activity | 3 | 1 | 0.004 |
| **GO:0099183** | BP | trans-synaptic signaling by BDNF, modulating synaptic transmission | 3 | 1 | 0.004 |
| **GO:0044090** | BP | positive regulation of vacuole organization | 2 | 1 | 0.005 |
| **GO:0030070** | BP | insulin processing | 2 | 1 | 0.005 |
| **GO:0043121** | MF | neurotrophin binding | 2 | 1 | 0.005 |
| **GO:1990667** | CC | PCSK9-AnxA2 complex | 2 | 1 | 0.006 |
| **GO:0045322** | MF | unmethylated CpG binding | 3 | 1 | 0.006 |
| **GO:0032804** | BP | negative regulation of low-density lipoprotein particle receptor catabolic process | 2 | 1 | 0.006 |
| **GO:0071593** | BP | lymphocyte aggregation | 2 | 1 | 0.006 |
| **GO:0016671** | MF | oxidoreductase activity, acting on a sulfur group of donors, disulfide as acceptor | 3 | 1 | 0.006 |
| **GO:1905581** | BP | positive regulation of low-density lipoprotein particle clearance | 2 | 1 | 0.006 |

| | | | | | |
|---|---|---|---|---|---|
| **GO:1905602** | BP | positive regulation of receptor-mediated endocytosis involved in cholesterol transport | 2 | 1 | 0.006 |
| **GO:1901895** | BP | negative regulation of calcium-transporting ATPase activity | 3 | 1 | 0.007 |
| **GO:0042997** | BP | negative regulation of Golgi to plasma membrane protein transport | 3 | 1 | 0.007 |
| **GO:0060263** | BP | regulation of respiratory burst | 4 | 1 | 0.007 |
| **GO:0045454** | BP | cell redox homeostasis | 69 | 2 | 0.007 |
| **GO:0045359** | BP | positive regulation of interferon-beta biosynthetic process | 7 | 1 | 0.007 |
| **GO:0032741** | BP | positive regulation of interleukin-18 production | 5 | 1 | 0.008 |
| **GO:0032640** | BP | tumor necrosis factor production | 4 | 1 | 0.008 |
| **GO:0034123** | BP | positive regulation of toll-like receptor signaling pathway | 4 | 1 | 0.008 |
| **GO:0048403** | MF | brain-derived neurotrophic factor binding | 3 | 1 | 0.008 |
| **GO:0098772** | MF | molecular function regulator | 3 | 1 | 0.009 |
| **GO:0050871** | BP | positive regulation of B cell activation | 5 | 1 | 0.009 |
| **GO:0002639** | BP | positive regulation of immunoglobulin production | 6 | 1 | 0.010 |
| **GO:0031340** | BP | positive regulation of vesicle fusion | 6 | 1 | 0.011 |
| **GO:0010310** | BP | regulation of hydrogen peroxide metabolic process | 6 | 1 | 0.011 |
| **GO:0031547** | BP | brain-derived neurotrophic factor receptor signaling pathway | 5 | 1 | 0.012 |

| | | | | | |
|---|---|---|---|---|---|
| **GO:0008330** | MF | protein tyrosine/threonine phosphatase activity | 4 | 1 | 0.012 |
| **GO:0002237** | BP | response to molecule of bacterial origin | 8 | 1 | 0.012 |
| **GO:0055069** | BP | zinc ion homeostasis | 5 | 1 | 0.012 |
| **GO:0097066** | BP | response to thyroid hormone | 6 | 1 | 0.012 |
| **GO:0032717** | BP | negative regulation of interleukin-8 production | 9 | 1 | 0.013 |
| **GO:0008142** | MF | oxysterol binding | 5 | 1 | 0.013 |
| **GO:0042490** | BP | mechanoreceptor differentiation | 5 | 1 | 0.015 |
| **GO:0050707** | BP | regulation of cytokine secretion | 6 | 1 | 0.015 |
| **GO:0035095** | BP | behavioral response to nicotine | 7 | 1 | 0.015 |
| **GO:0035197** | MF | siRNA binding | 9 | 1 | 0.015 |
| **GO:0046548** | BP | retinal rod cell development | 8 | 1 | 0.015 |
| **GO:0004725** | MF | protein tyrosine phosphatase activity | 83 | 2 | 0.015 |
| **GO:0032460** | BP | negative regulation of protein oligomerization | 7 | 1 | 0.016 |
| **GO:0005149** | MF | interleukin-1 receptor binding | 9 | 1 | 0.016 |
| **GO:0042129** | BP | regulation of T cell proliferation | 10 | 1 | 0.016 |
| **GO:0010989** | BP | negative regulation of low-density lipoprotein particle clearance | 16 | 1 | 0.016 |
| **GO:0022417** | BP | protein maturation by protein folding | 9 | 1 | 0.017 |
| **GO:0035749** | CC | myelin sheath adaxonal region | 6 | 1 | 0.017 |
| **GO:0043304** | BP | regulation of mast cell degranulation | 8 | 1 | 0.017 |
| **GO:0036035** | BP | osteoclast development | 5 | 1 | 0.017 |
| **GO:0046486** | BP | glycerolipid metabolic process | 5 | 1 | 0.017 |

| | | | | | |
|---|---|---|---|---|---|
| **GO:0045078** | BP | positive regulation of interferon-gamma biosynthetic process | 12 | 1 | 0.017 |
| **GO:0007589** | BP | body fluid secretion | 9 | 1 | 0.018 |
| **GO:0001765** | BP | membrane raft assembly | 6 | 1 | 0.018 |
| **GO:0004713** | MF | protein tyrosine kinase activity | 78 | 2 | 0.018 |
| **GO:0070286** | BP | axonemal dynein complex assembly | 9 | 1 | 0.018 |
| **GO:0032725** | BP | positive regulation of granulocyte macrophage colony-stimulating factor production | 11 | 1 | 0.018 |
| **GO:0019887** | MF | protein kinase regulator activity | 7 | 1 | 0.019 |
| **GO:0015464** | MF | acetylcholine receptor activity | 10 | 1 | 0.019 |
| **GO:0005975** | BP | carbohydrate metabolic process | 118 | 2 | 0.019 |
| **GO:0097371** | MF | MDM2/MDM4 family protein binding | 8 | 1 | 0.019 |
| **GO:0022849** | MF | glutamate-gated calcium ion channel activity | 5 | 1 | 0.020 |
| **GO:1903997** | BP | positive regulation of non-membrane spanning protein tyrosine kinase activity | 7 | 1 | 0.020 |
| **GO:0006900** | BP | vesicle budding from membrane | 9 | 1 | 0.020 |
| **GO:0032009** | CC | early phagosome | 11 | 1 | 0.021 |
| **GO:0005892** | CC | acetylcholine-gated channel complex | 11 | 1 | 0.021 |
| **GO:0019227** | BP | neuronal action potential propagation | 9 | 1 | 0.021 |
| **GO:0043410** | BP | positive regulation of MAPK cascade | 95 | 2 | 0.022 |
| **GO:0007252** | BP | I-kappaB phosphorylation | 12 | 1 | 0.022 |
| **GO:0035335** | BP | peptidyl-tyrosine dephosphorylation | 99 | 2 | 0.022 |
| **GO:0000791** | CC | Euchromatin | 8 | 1 | 0.022 |

| | | | | | |
|---|---|---|---|---|---|
| **GO:0044354** | CC | Macropinosome | 8 | 1 | 0.022 |
| **GO:0005671** | CC | Ada2/Gcn5/Ada3 transcription activator complex | 15 | 1 | 0.023 |
| **GO:0004970** | MF | ionotropic glutamate receptor activity | 7 | 1 | 0.024 |
| **GO:0005858** | CC | axonemal dynein complex | 10 | 1 | 0.025 |
| **GO:0043220** | CC | Schmidt-Lanterman incisure | 11 | 1 | 0.025 |
| **GO:0016558** | BP | protein import into peroxisome matrix | 10 | 1 | 0.025 |
| **GO:0034162** | BP | toll-like receptor 9 signaling pathway | 16 | 1 | 0.025 |
| **GO:0045577** | BP | regulation of B cell differentiation | 11 | 1 | 0.025 |
| **GO:0004972** | MF | NMDA glutamate receptor activity | 7 | 1 | 0.025 |
| **GO:0006893** | BP | Golgi to plasma membrane transport | 13 | 1 | 0.026 |
| **GO:0045647** | BP | negative regulation of erythrocyte differentiation | 10 | 1 | 0.026 |
| **GO:0016580** | CC | Sin3 complex | 12 | 1 | 0.026 |
| **GO:0060285** | BP | cilium-dependent cell motility | 11 | 1 | 0.026 |
| **GO:0036020** | CC | endolysosome membrane | 13 | 1 | 0.027 |
| **GO:0045779** | BP | negative regulation of bone resorption | 12 | 1 | 0.027 |
| **GO:0021954** | BP | central nervous system neuron development | 13 | 1 | 0.027 |
| **GO:0097553** | BP | calcium ion transmembrane import into cytosol | 7 | 1 | 0.028 |
| **GO:0015276** | MF | ligand-gated ion channel activity | 13 | 1 | 0.028 |
| **GO:0051770** | BP | positive regulation of nitric-oxide synthase biosynthetic process | 14 | 1 | 0.028 |
| **GO:0030277** | BP | maintenance of gastrointestinal epithelium | 11 | 1 | 0.028 |

| | | | | | |
|---|---|---|---|---|---|
| **GO:0022848** | MF | acetylcholine-gated cation-selective channel activity | 16 | 1 | 0.028 |
| **GO:0005779** | CC | integral component of peroxisomal membrane | 16 | 1 | 0.029 |
| **GO:0043968** | BP | histone H2A acetylation | 13 | 1 | 0.030 |
| **GO:0098976** | BP | excitatory chemical synaptic transmission | 8 | 1 | 0.030 |
| **GO:0050765** | BP | negative regulation of phagocytosis | 16 | 1 | 0.030 |
| **GO:0033673** | BP | negative regulation of kinase activity | 13 | 1 | 0.031 |
| **GO:0048935** | BP | peripheral nervous system neuron development | 10 | 1 | 0.031 |
| **GO:0032722** | BP | positive regulation of chemokine production | 19 | 1 | 0.031 |
| **GO:0004143** | MF | diacylglycerol kinase activity | 11 | 1 | 0.031 |
| **GO:0035267** | CC | NuA4 histone acetyltransferase complex | 14 | 1 | 0.031 |
| **GO:0098691** | CC | dopaminergic synapse | 14 | 1 | 0.033 |
| **GO:0005548** | MF | phospholipid transporter activity | 14 | 1 | 0.033 |
| **GO:0034122** | BP | negative regulation of toll-like receptor signaling pathway | 17 | 1 | 0.033 |
| **GO:0006470** | BP | protein dephosphorylation | 127 | 2 | 0.033 |
| **GO:0032733** | BP | positive regulation of interleukin-10 production | 25 | 1 | 0.035 |
| **GO:0031982** | CC | Vesicle | 130 | 2 | 0.035 |
| **GO:0036150** | BP | phosphatidylserine acyl-chain remodeling | 21 | 1 | 0.035 |
| **GO:1902041** | BP | regulation of extrinsic apoptotic signaling pathway via death domain receptors | 17 | 1 | 0.035 |
| **GO:0001964** | BP | startle response | 13 | 1 | 0.035 |

| | | | | | |
|---|---|---|---|---|---|
| **GO:0003887** | MF | DNA-directed DNA polymerase activity | 21 | 1 | 0.035 |
| **GO:0046834** | BP | lipid phosphorylation | 12 | 1 | 0.035 |
| **GO:0007031** | BP | peroxisome organization | 19 | 1 | 0.036 |
| **GO:0034236** | MF | protein kinase A catalytic subunit binding | 13 | 1 | 0.036 |
| **GO:0030659** | CC | cytoplasmic vesicle membrane | 133 | 2 | 0.036 |
| **GO:0033198** | BP | response to ATP | 15 | 1 | 0.037 |
| **GO:0044548** | MF | S100 protein binding | 14 | 1 | 0.038 |
| **GO:0010243** | BP | response to organonitrogen compound | 16 | 1 | 0.038 |
| **GO:0043235** | CC | receptor complex | 127 | 2 | 0.039 |
| **GO:0090023** | BP | positive regulation of neutrophil chemotaxis | 24 | 1 | 0.039 |
| **GO:0017146** | CC | NMDA selective glutamate receptor complex | 11 | 1 | 0.039 |
| **GO:0045859** | BP | regulation of protein kinase activity | 20 | 1 | 0.039 |
| **GO:0001921** | BP | positive regulation of receptor recycling | 14 | 1 | 0.039 |
| **GO:0051019** | MF | mitogen-activated protein kinase binding | 18 | 1 | 0.040 |
| **GO:0007631** | BP | feeding behavior | 23 | 1 | 0.040 |
| **GO:0098641** | MF | cadherin binding involved in cell-cell adhesion | 17 | 1 | 0.042 |
| **GO:0018108** | BP | peptidyl-tyrosine phosphorylation | 120 | 2 | 0.042 |
| **GO:2000811** | BP | negative regulation of anoikis | 17 | 1 | 0.042 |
| **GO:0071375** | BP | cellular response to peptide hormone stimulus | 16 | 1 | 0.043 |
| **GO:0006259** | BP | DNA metabolic process | 23 | 1 | 0.043 |
| **GO:0010592** | BP | positive regulation of lamellipodium assembly | 20 | 1 | 0.043 |

| | | | | | |
|---|---|---|---|---|---|
| **GO:0019897** | CC | extrinsic component of plasma membrane | 21 | 1 | 0.044 |
| **GO:0032735** | BP | positive regulation of interleukin-12 production | 25 | 1 | 0.044 |
| **GO:0010039** | BP | response to iron ion | 18 | 1 | 0.044 |
| **GO:0060044** | BP | negative regulation of cardiac muscle cell proliferation | 21 | 1 | 0.044 |
| **GO:0003951** | MF | NAD+ kinase activity | 17 | 1 | 0.045 |
| **GO:0002755** | BP | MyD88-dependent toll-like receptor signaling pathway | 32 | 1 | 0.046 |
| **GO:0022011** | BP | myelination in peripheral nervous system | 17 | 1 | 0.047 |
| **GO:0070064** | MF | proline-rich region binding | 18 | 1 | 0.047 |
| **GO:0016575** | BP | histone deacetylation | 22 | 1 | 0.047 |
| **GO:0006298** | BP | mismatch repair | 22 | 1 | 0.047 |
| **GO:0030301** | BP | cholesterol transport | 21 | 1 | 0.048 |
| **GO:0035902** | BP | response to immobilization stress | 23 | 1 | 0.049 |
| **GO:0032728** | BP | positive regulation of interferon-beta production | 26 | 1 | 0.050 |
| **GO:0045453** | BP | bone resorption | 22 | 1 | 0.050 |
| **GO:0048011** | BP | neurotrophin TRK receptor signaling pathway | 20 | 1 | 0.050 |

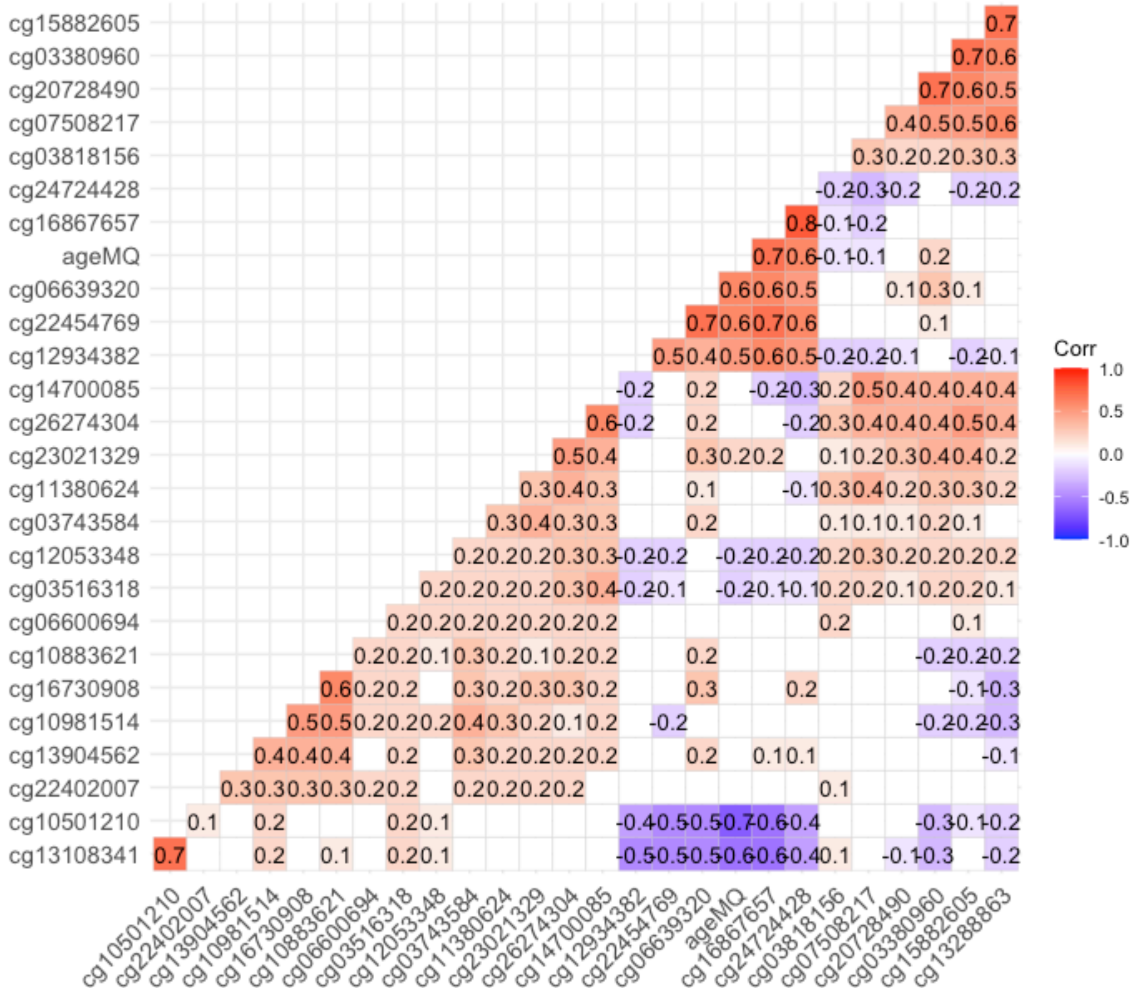


Supplementary Figure 1: Correlations of dmCpGs related to fathers' smoking onset before age 15 years with aging marker CpGs from the RHINESSA Cohort (cg1686765: ELOVL2, cg24724428: ELOVL2, cg22454769: FHL2 and cg131083: DNAH9) and Offspring age. 9 dmCpGs showed correlation = 0 with age; aging methylation markers correlation >=|0.6|.

Supplementary Table E10: Associations between dmCpGs identified in relation to fathers' smoking onset before age 15years and phenotypic outcomes in the offspring (adjusted for offspring's sex).

| Offspring outcomes | CpG site | Coefficient | P value | Gene |
|---|---|---|---|---|
| Ever asthma | cg22402007 | 10.7900 | 0.014 | NTRK2 |
| Ever wheezing | cg11380624 | 14.4008 | 0.000 | DNAJC14 |
| Ever wheezing | cg10981514 | -8.3351 | 0.012 | TPCN1 |
| Weight | cg12053348 | -51.0810 | 0.001 | Intergenic |
| Weight | cg03380960 | 42.5750 | 0.023 | FAM53B |
| Weight | cg22402007 | -67.1670 | 0.013 | NTRK2 |
| BMI | cg23021329 | 0.0024 | 0.067 | TLR9 |
| BMI | cg12053348 | -0.0013 | 0.004 | Intergenic |
| BMI | cg03380960 | 0.0022 | 0.008 | FAM53B |
| BMI | cg22402007 | -0.0017 | 0.025 | NTRK2 |
| BMI | cg11380624 | -0.0013 | 0.074 | DNAJC14 |

Supplementary Table E11: Power calculations using R package pwrEWAS[28]

| Samples | Target Delta | | | |
|---|---|---|---|---|
| | 0.05 | 0.10 | 0.15 | 0.20 |
| A: preconception father smoking: effect size=0.024, sample split=0.62, target dmCpGs=4, FDR=0.05, tissue type=Blood adult | | | | |
| 500 | 0.79 | 0.88 | 0.92 | 0.95 |
| 600 | 0.85 | 0.92 | 0.95 | 0.97 |
| 700 | 0.89 | 0.95 | 0.97 | 0.98 |
| 800 | 0.93 | 0.97 | 0.98 | 0.99 |
| 900 | 0.95 | 0.98 | 0.99 | 0.99 |
| B: Start smoking before age <15; effect size=0.02, sample split=0.21, target dmCpGs=19, FDR=0.05, tissue type=Blood adult | | | | |
| 200 | 0.64 | 0.76 | 0.83 | 0.88 |
| 250 | 0.72 | 0.83 | 0.88 | 0.92 |
| 300 | 0.79 | 0.88 | 0.92 | 0.95 |
| 350 | 0.84 | 0.91 | 0.95 | 0.96 |
| 400 | 0.88 | 0.94 | 0.96 | 0.98 |
